# Innate detection of *Salmonella* replication triggers caspase-8-dependent apoptosis via TLR-driven TNF signaling and NLRC4-mediated sensing of the SPI-2 Type III secretion system

**DOI:** 10.1101/2025.09.10.675376

**Authors:** Beatrice I. Herrmann, Maxime Zamba-Campero, Reyna Garcia-Sillas, Winslow W. Yost, Lance W. Peterson, Justin L. Roncaioili, Shelley C. Rankin, Dieter M. Schifferli, Kodi Ravichandran, Igor E. Brodsky

## Abstract

*Salmonella enterica* comprises over 2500 serovars that are responsible for over 90 million annual infections and 100,000 deaths worldwide. Despite this diversity, our understanding of innate immune responses to *Salmonella* is based on extensive study of a few serovars, primarily Typhimurium, including strains that cannot replicate within primary murine macrophages. Non-replicating *Salmonella* trigger caspase-1 and −11-dependent pyroptosis. Whether the innate immune system distinguishes between replicating and non-replicating intracellular *Salmonella* is poorly defined. Here we demonstrate that replicating *Salmonella enterica* induce a distinct pathway of TNF- and caspase-8-driven apoptosis via host TLR4 and *Salmonella* Pathogenicity Island-2 activity. This pathway is independent of gasdermin D and involves the apoptotic pore protein Pannexin-1. Combined loss of Pannexin-1 and gasdermin D resulted in defective control of systemic *Salmonella*, indicating that these pathways function together to promote anti-*Salmonella* host defense. Altogether, our findings uncover a previously unappreciated pathway by which macrophages sense intracellular replicating bacteria.

## Introduction

*Salmonella enterica* is a gram-negative intracellular pathogen that replicates within mammalian cells and causes severe disease ranging from self-limiting gastroenteritis to systemic bacteremia and chronic infection. *Salmonella* infection constitutes a major global health burden that is responsible for over 90 million enteric infections and 100,000 deaths annually worldwide due to diarrheal disease, with the majority of global human infections caused by the serovars Typhimurium and Enteritidis^1,2^.

All *Salmonella enterica* encode two type III secretion systems (T3SS) on *Salmonella* pathogenicity islands (SPI-1 and SPI-2), that respectively enable *Salmonella* to invade non-phagocytic cells, and survive in immune cells within the *Salmonella-*containing vacuole (SCV)^3,4^. However, innate immune sensing of *Salmonella* SPI-1 T3SS activity triggers pyroptosis that mediate host defense and control *Salmonella* infection^5–12^.

Pyroptosis is a lytic, inflammatory cell death program that is mediated by the pore-forming protein gasdermin D (GSDMD) following its cleavage by caspases-1 and/or −11 (Casp1/11)^13–17,27,30^. Pyroptosis controls bacterial invasion and spread by eliminating infected cells and promoting release of IL-1 family cytokines^11,12,18^. During the intestinal stage of infection, *Salmonella* expresses flagella and the SPI-1 T3SS, which enables *Salmonella* to invade intestinal epithelial cells and can be recapitulated *in vitro* by growth of the bacteria in the presence of high salt and low oxygen^19,20^. Macrophages that encounter invasive *Salmonella* undergo rapid Casp1 activation via the NLR family of apoptosis inhibitory proteins (NAIP)/NLRC4 inflammasome that detects cytosolic flagellin and SPI-1 T3SS needle and rod proteins^7,21–24,14,25^. In contrast, non-SPI-1-expressing *Salmonella* are phagocytosed and are detected at later stages of infection via sensing of cytosolic bacterial lipopolysaccharide (LPS)^16,26–28^. Cytosolic LPS triggers delayed activation of the noncanonical Casp11 inflammasome and subsequent pyroptosis independently of the canonical NAIP/NLRC4 inflammasome and requires the activity of the SPI-2 T3SS^17,27–29^. Notably, if the ability of the cell to undergo pyroptosis is disrupted, apoptosis can mediate cell death in response to intracellular bacteria, highlighting the importance of cell death in antibacterial defense^30–33^. In contrast to pyroptosis, apoptosis is classically viewed as a non-inflammatory programmed cell death driven by the initiator caspases-8 (Casp8) or −9 and comprises an ordered process of cellular disassembly that maintains membrane integrity and promotes clearance of dying cells via efferocytosis^28,34–40^. In the absence of Casp1, Casp8 can associate with the NAIP/NLRC4 inflammasome to mediate a GSDMD-independent cell death in response to detection of bacterial flagellin or SPI-1 T3SS components ^33,41–46^.

Our understanding of innate responses to *Salmonella* have largely been based on studies using a limited number of *Salmonella* serovars, including the *S*. Typhimurium strain SL1344. Although it is virulent in rodent infection models, SL1344 cannot replicate in primary murine macrophages due to mutation of the *hisG* gene, and its associated histidine auxotrophy^47–49^. These foundational studies established that macrophages detect intracellular *Salmonella* primarily via the Casp11 non-canonical inflammasome^18,29^. These findings raise the question of whether the innate immune system detects replicating intracellular pathogens and distinguishes them from viable non-replicating organisms. Pattern recognition receptor signaling upregulates inflammatory gene expression programs, and live intracellular bacteria are detected via the presence of molecules known as Vita-PAMPs, which trigger particular pathways of inflammasome activation^50–52^. However, whether the innate immune system responds differently to replicating bacteria compared to similar viable non-replicating bacterial pathogens is not clear.

Here we demonstrate that intracellular replicating *Salmonella*, including the globally prevalent serovar *S*. Enteritidis, invasive non-typhoidal *S*. Typhimurium, and a histidine prototrophic revertant of SL1344, all trigger a previously undescribed Casp1/11-*independent* apoptosis that is mediated by caspase-8 (Casp8). This response occurs in the presence of Casp1/11, indicating that it is not solely a backup pathway, though it is amplified under Casp1/11-deficiency. Loss of histidine prototrophy or disruption of SPI-2 function abrogates both bacterial replication and Casp8-mediated apoptosis, which is GSDMD-independent and involves cleavage of the apoptotic pore Pannexin1. Mechanistically, elevated bacterial replication results in increased TLR4-dependent production of the inflammatory cytokine TNF, and TLR4 and TNFR1 signaling are required for Casp8-dependent apoptosis in response to *Salmonella* replication. Interestingly, Casp8 also participates in a second apoptosis pathway downstream of *Salmonella* replication that involves NLRC4-dependent sensing of SPI-2 T3SS structural components that are present at elevated levels in the setting of intracellular replication. Importantly, loss of Casp8-dependent apoptosis results in decreased survival and increased bacterial burdens *in vivo*. Altogether, these findings reveal that replicating intracellular *Salmonella* trigger a distinct pathway of Casp8-dependent apoptosis that promotes anti-bacterial immune defense.

## Results

### Caspase-8 is a non-redundant mediator of innate control of *Salmonella* infection

Wild-type C57B6L (WT) mice undergo an acute lethal infection with *Salmonella enterica* serovar Typhimurium (*S*.Tm)^11,53^. The SL1344 strain of S.Tm has been extensively used in both mechanistic *in vitro* studies of innate immune responses to *Salmonella*, as well as *in vivo* studies of *Salmonella* pathogenesis^47,48^. We sought to extend our understanding of *Salmonella-*host interactions by investigating immune responses to other serovars of *Salmonella enterica* as well as other strains of *S*. Typhimurium. Interestingly, in contrast to SL1344 (*S*.Tm), we found that *S. enterica* serovar Enteritidis (*S*.Ent) isolate obtained from the University of Pennsylvania School of Veterinary Medicine *Salmonella* strain collection did not induce lethality following intraperitoneal infection in WT mice or mice lacking caspases-1 and −11 (*Casp1/11^−/−^*) (Fig. 1a), similar to reports of some human clinical isolates of *S*. Enteritidis that are also non-lethal in C57BL6 mice^54^. Notably, *S*. Ent persisted in tissues of both WT and *Casp1/11^−/−^* at least to 60 days post-infection, with *Casp1/11*^−/−^ mice having significantly higher bacterial tissue burdens on day 14 post-infection (Fig. 1b-d). These data indicate that *S*.Ent can establish long term carriage in B6 mice, and that while Casp1/11 contribute to early control of *S.*Ent, they are not absolutely required for survival of WT *Salmonella* systemic infection.

**Fig. 1.**
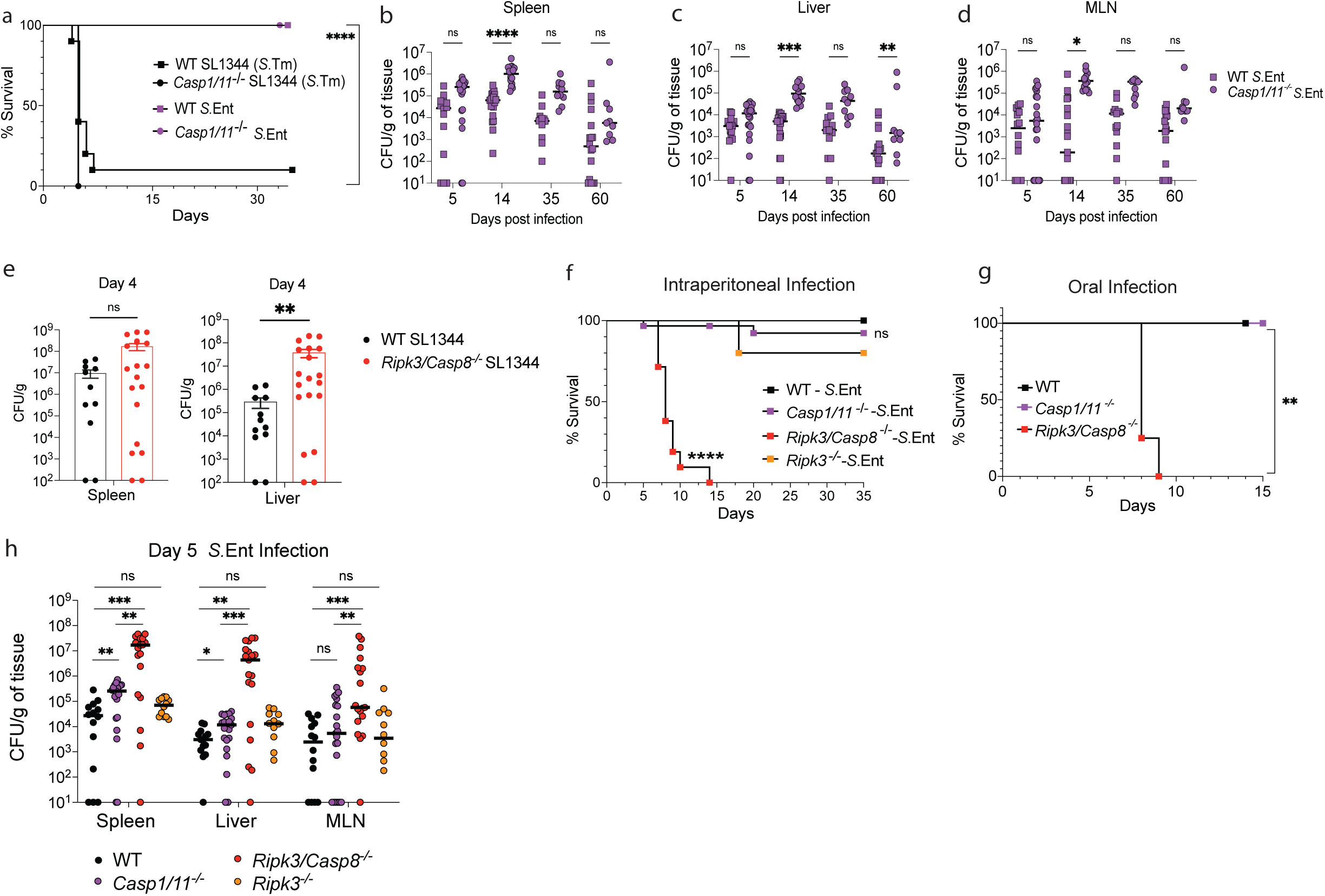
Caspase-8 is a non-redundant mediator of innate control of *Salmonella* infection. **a-d,** Groups of B6 mice were infected with 1×10^3^ CFU of indicated *Salmonella* strains via intraperitoneal injection. **a,** Survival of WT and *Casp1/11^−/−^* mice infected with SL1344 (n=10, n=9) or *S.*Ent 2015-3 (n=8,n=8). **b-d,** *S.*Ent bacterial burdens (CFU g^−1^ tissue) in the spleen, liver, and mesenteric lymph-nodes at 5-, 14-, 35-, and 60-days post-infection. **e,** SL1344 bacterial burdens (CFU g^−1^ tissue) in spleen and liver of WT and *Ripk3/Casp8^−/−^* mice at 4 days post i.p. infection **f,** Survival curves of WT (n=21), *Casp1/11^−/−^* (n=30), *Ripk3*^−/−^ (n=5), *Ripk3/Casp8^−/−^* (n=21) infected with *S.*Ent over 35 days. **g,** WT (n=5), *Casp1/11^−/−^* (n=5), *Ripk3/Casp8^−/−^* (n=4) infected with *S.*Ent (1×10^8^ CFU) via oral gavage. **h,** *S.*Ent bacterial burdens (CFU g^−1^ tissue) in the spleen, liver, and mesenteric lymph-nodes at 5 days post-infection. Kaplan Meier curves for survival statistics. Mann–Whitney *U* test (two-tailed) for bacterial burdens. \**P* < 0.05, \*\**P* < 0.01, \*\*\*\**P* < 0.0001; NS, not significant.

Casp1/11 are thought to be the primary caspases that mediate cell death and host protection in response to *Salmonella* infection^14,18,28^ with Casp8 serving as a backup when Casp1/11 are missing^33^. Studies examining the role of caspases to control of *Salmonella* infection concluded that other caspases are engaged when Casp1/11 were absent, and particularly that Casp1/11/12/8 collectively compensate for one another’s loss to mediate anti-*Salmonella* host defense^33^. However, these studies employed multi-auxotrophic *S*.Tm strains that lack both histidine and aromatic amino acid biosynthesis (SL1344 Δ*aroA*). Whether Casp8 plays a distinct, non-redundant role in host defense against acute *Salmonella* infection has not been tested. The study of murine Casp8 requires concomitant deletion of *Ripk3*, to avoid embryonic lethality due to RIPK3-dependent necroptosis^55–57^. Notably, while previous reports indicated no significant defects in control of SL1344 Δ*aroA* burdens in mice lacking *Ripk3/*Casp8, we found these mice harbored significantly higher SL1344 burdens as early as day 4 post-infection (Fig. 1e). Furthermore, both *Ripk3*^−/−^ and *Casp1/11^−/−^* mice had no significant defect in survival of *S.*Ent infection, whereas *Ripk3/Casp8*^−/−^ mice all succumbed to infection within 14 days of either intraperitoneal or oral infection (Fig. 1f, g). *Ripk3/Casp8^−/−^* mice also had significantly higher *S*.Ent tissue burdens compared to *Casp1/11^−/−^*, *Ripk3^−/−^*, and WT mice five days post-infection (Fig. 1h). Altogether, these data demonstrate that Casp8 plays a nonredundant role in control of wild-type systemic *S. enterica in vivo*, raising questions about how Casp8 contributes to *Salmonella-*induced macrophage cell death during systemic infection.

### *Salmonella* Enteritidis activates caspase-8- and RIPK1 kinase-dependent cell death independently of caspase-1 and −11

Under systemic infection conditions when the SPI-1 T3SS is not thought to be highly expressed, macrophages phagocytose *Salmonella,* which traffics to the *Salmonella-*containing vacuole (SCV) where it upregulates the SPI-2 T3SS which promotes bacterial survival and replication^58^. infected under non-invasive conditions, when *Salmonella* enters the cell via phagocytosis, this manner exhibit delayed cell death 16-20 hours post-infection via activation of Casp11 in response to bacterial LPS, and subsequent NLRP3/Casp1 inflammasome activation downstream of Casp11-dependent GSDMD pore formation^18,29^. This contrasts with rapid NAIP/NLRC4 inflammasome activation triggered by cytosolic flagellin, SPI-1 needle, or rod proteins that are expressed under conditions that mimic the intestinal stage of infection^14,23,25^. Casp8 compensates for Casp1 deficiency during invasive *Salmonella*-infection^33,43,46^. Under non-invasive conditions however, SL1344 *S*.Tm triggers Casp1/11-dependent delayed pyroptosis (Fig. 2a)^18,28,29^. Unexpectedly, *S*.Ent-induced cell death was only slightly reduced in *Casp1/11^−/−^* macrophages and was abrogated in macrophages lacking *Casp1/11/Ripk3/Casp8* (Fig. 2a). RIPK3-dependent programmed necrosis was not responsible for this, as cell death in isogenic *Casp8^+/−^* macrophages did not differ from *Casp1/11^−/−^* macrophages, and sole RIPK3 deficiency or RIPK3 inhibitor treatment had no effect (Fig. 2a, Extended Data Fig. 1a,b). Critically, multiple other *S. enterica* strains and serovars, including the commonly used *S*.Tm strain (ATCC14028), invasive non-typhoidal *S*.Tm (DT104), another *S*.Ent isolate and *S*. Dublin isolate both from the PennVet strain collection, all induced Casp1/11-independent, Casp8-dependent cell death (Extended Data Fig. 1c), indicating that this is *S. enterica* generally induces Casp1/11- and SPI-1-independent macrophage cell death that requires Casp8. These strains all triggered similar levels of Casp1-dependent macrophage cytotoxicity to SL1344 under SPI-1-inducing conditions, indicating that Casp8-dependent late cell death is distinct from rapid NLRC4/NAIP-dependent pyroptosis mediated by recognition of *Salmonella* flagellin and SPI-1 (Extended Data Fig. 1d). Consistently, deletion of *S*.Ent flagellin did not impact Casp1/11-independent cell death (Extended Data Fig. 1e).

**Fig. 2.**
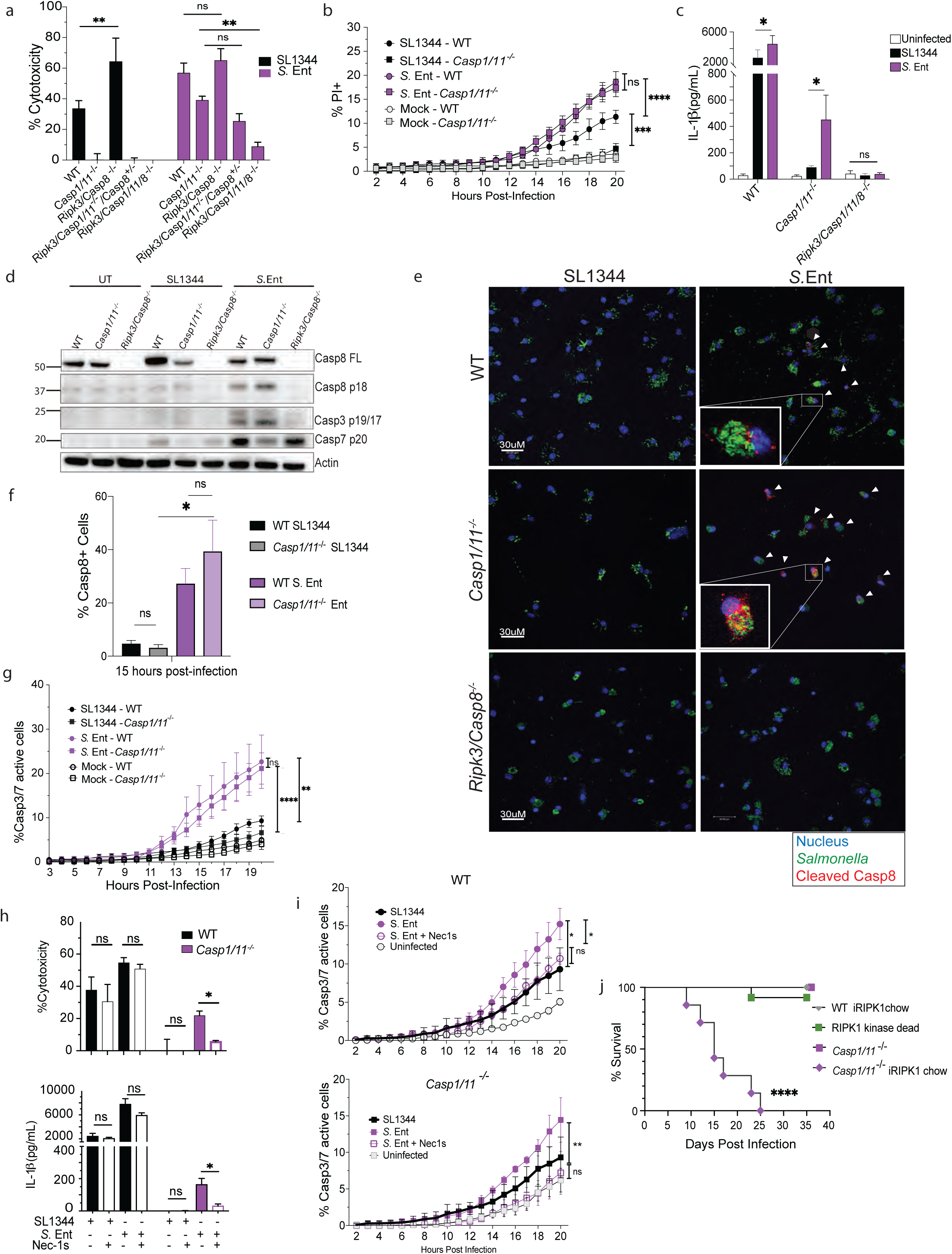
*Salmonella* Enteritidis activates caspase-8- and RIPK1 kinase-dependent cell death independently of caspase-1 and −11. **a,** Cytotoxicity determined by LDH release in primary WT, *Casp1/11^−/−^*, *Ripk3/Casp8^−/−^*, *Ripk3/Casp1/11^−/−^Casp8^+/−^*, *Ripk3/Casp1/11/8^−/−^* BMDMs infected at MOI of 100 for 18 hours with SL1344 or *S.*Ent. **b,** Infected cells stained with propidium iodide (PI) were placed within the IncuCyte live-cell imaging analysis system. Percent propidium iodide uptake measured by analysis of PI^+^ cells versus total cells counted at indicated timepoints. Statistical comparisons between cell death kinetics of SL1344 and *S.*Ent and indicated genotypes. **c,** IL-1β cytokine release from BMDMs determined by Cytometric Bead Array. **d,** Immuno-blot analysis of caspase cleavage in combined whole cell lysates and supernatants of primary BMDMs harvested 18 hours post-infection. β-actin is used as loading control. **e,** Confocal microscopy of WT, *Casp1/11^−/−^*, *Ripk3/Casp8^−/−^* BMDMs infected for 15 hours with indicated SL1344 and *S.*Ent strains expressing GFP. Cleaved Casp8 (red) and nuclear Hoechst 33342 counterstain (blue). White arrows indicate cells containing cleaved Casp8. **f,** Quantification of cells positive for Casp8 cleavage. **g,** Casp3/7 IncuCyte kinetic assay of WT and *Casp1/11^−/−^* BMDMs infected with either SL1344 or *S.*Ent. Biological replicate representative of 3 experiments. **h,** LDH release and IL-1β release in BMDMs infected with SL1344 or *S*.Ent and treated with Nec-1s. **i,** Casp3/7 IncuCyte kinetic assay of WT or *Casp1/11^−/−^* BMDMs infected with SL1344 or *S*.Ent and either mock or Nec-1s treated. Percentage of Casp3/7 active cells tracked over the course of infection. Data represent mean ± SEM of at least 3 independent experiments. \**P*<0.05 \*\**P*<0.01 \*\*\**P*<0.001 by One-way ANOVA (a,c,f,h,i) or Two-way ANOVA (b,g,i). **j,** B6 mice were placed on a diet of irradiated Purina 5001 control chow or chow containing RIPK1 inhibitor GSK3540547A (GSKʹ547 chow, provided by GSK) 3 days prior to infection and allowed to eat ad libitum for the duration of the experiment. Survival ratio of WT fed inhibitor chow (n=10), RIPK1^KD^ (n=12), or *Casp1/11^−/−^* fed control chow (n=7), or inhibitor chow (n=9). \**P* < 0.05, \*\**P* < 0.01, \*\*\*\**P* < 0.0001 by (Two-tailed)-Whitney *U* test. NS, not significant.

Membrane permeability as measured by both propidum iodide or SYTOX staining in response to *Salmonella* infection began around 12 hours post-infection, highlighting the delayed nature of cell death under these conditions (Fig. 2b, Extended Data Fig. 1f). While SL1344-induced membrane permeability increased only modestly after 12 hours, *S.*Ent-induced membrane damage continued to increase and was significantly elevated relative to SL1344-infected cells by 17 hours post-infection (Fig. 2b, Extended Data Fig. 1f). Notably, S. Ent induced identical levels of membrane permeability in *Casp1/11^−/−^* and WT BMDMs, whereas SL1344 induced lower permeability in *Casp1/11^−/−^* BMDMs compared to WT BMDMs (Fig. 2b, Extended Data Fig. 1f). These data suggest that *S*.Ent induces Casp8-driven membrane permeability independently of Casp1/11.

*S.*Ent also induced increased processing and release of IL-1 cytokines compared to SL1344 in WT BMDMs (Fig. 2c, Extended Data Fig. 2g, h), and induced significant Casp1/11-independent IL-1 release, which was Casp8-dependent (Fig. 2c, Extended Data Fig. 2e,f). Levels of pro-IL-1β were similar in SL1344- and *S*.Ent-infected cells, and the inflammasome-independent cytokine IL-6 was not significantly elevated in *S.*Ent-infected BMDMs compared to those infected with SL1344, indicating that increased IL-1 release was not due to altered expression of pro-IL-1 in response to *S*.Ent (Extended Data Fig. 2g, i).

Casp8 cleaves and activates the apoptotic caspases-3 (Casp3) and −7 (Casp7)^34^. Consistently, *S.*Ent triggered Casp8, Casp3, and Casp7 processing in both WT and *Casp1/11*^−/−^ BMDMs, whereas SL1344 did not induce activation of Casp8 or Casp3 (Fig. 2d). Consistent with the known role of Casp1 in Casp7 processing^59^, we observed Casp7 processing in both WT and *Ripk3/Casp8^−/−^* BMDMs infected with SL1344 and *S*.Ent but not in SL1344-infected BMDMs lacking Casp1/11 (Fig. 2d). Consistently, immunofluorescence microscopy revealed cleaved Casp8 in *S*.Ent-infected WT BMDMs that was elevated in Casp1/11-deficient macrophages and virtually absent in SL1344-infected BMDMs (Fig. 2e,f). Furthermore, Casp3/7 activation occurred with similar kinetics and magnitude in WT and *Casp1/11*-deficient BMDMs following *S*.Ent infection (Fig. 2g). In contrast, SL1344-infected WT macrophages exhibited only minimal Casp3/7 activity, that was reduced to mock-infected levels in *Casp1/11^−/−^* BMDMs (Fig. 2g). Together these data demonstrate that *S.*Ent infection triggers Casp8 activation in both WT and *Casp1/11^−/−^* BMDMs, indicating that Casp8 activation during *S*.Ent infection is independent of Casp1/11 deficiency.

Casp8 activation in response to bacterial infection engages Receptor Interacting Protein Kinase 1 (RIPK1)^60–63^. However, use of Nec1s treatment alone to block RIPK1 kinase activity in WT cells^64^ did not affect cell death or IL-1β release, similar to *Ripk3Casp8^−/−^* BMDMs (Fig. 2h). However, inhibiting RIPK1 kinase activity in *Casp1/11^−/−^* cells completely abrogated *S*.Ent-triggered cell death and IL-1β release, and abrogated *Casp3/7* activity in *Casp1/11^−/−^* BMDMs (Fig. 2h, i). Critically, while chemically or genetically ablating RIPK1 kinase activity *in vivo*^65^ did not affect host survival following *S.*Ent infection, *Casp1/11*-deficient mice treated with RIPK1 inhibitor chow exhibited a profound defect in survival following oral infection (Fig. 2j). Altogether, these findings demonstrate that *Salmonella enterica* infection activates Casp8 independently of *Casp1/11*-deficiency and suggests that Casp8-induced cell death promotes host defense against wild-type *Salmonella in vivo*.

### Pannexin-1 promotes Casp8-dependent cell death in response to *Salmonella* infection

In addition to initiating extrinsic apoptosis, Casp8 can also mediate pyroptosis by cleaving GSDMD and promoting IL-1β release in response to *Yersinia* infection^63,66,67,68,69^. However, although GSDMD was cleaved to its active pore forming fragment (p30) in both SL1344 and *S.*Ent-infected WT cells, this cleavage was abrogated in *Casp1/11^−/−^* but not *Ripk3/Casp8^−/−^* BMDMs, indicating that Casp8 does not activate GSDMD during intracellular *Salmonella* infection (Fig. 3a). In contrast, we observed the presence of the N-terminal GSDMD p20 fragment, which is an indicator of GSDMD *inactivation* by Casp3^62,70^, in *S*.Ent-infected cell lysates in WT and *Casp1/11^−/−^* BMDMs, but not Casp8-deficient BMDMs, indicating that Casp8 is necessary for GSDMD inactivation, likely via Casp3 cleavage (Fig. 3a, Extended Data Fig. 2a). Consistently, GSDME, which is cleaved and activated by Casp3^71,62,69^, was cleaved in *S*.Ent-infected cells in a Casp8-dependent manner (Fig 3a, Extended Data Fig. 2b), consistent with robust Casp3/7 activity in *S*.Ent-infected cells. Importantly, while Casp1/11 and GSDMD both contributed to cell death in *S*.Ent-infected cells, combined deletion of Casp1/11 and GSDMD did not further reduce this cytotoxicity, indicating that Casp1/11 and GSDMD mediate cell death via the same pathway (Fig. 3b). Notably, GSDME was not responsible for Casp1/11/GSDMD-independent cell death, as either deletion of GSDME alone or in cells lacking GSDMD did not impact cell death (Fig. 3b). Critically however, loss of both GSDMD and Casp8, significantly reduced cell death following *S*.Ent infection (Fig. 3c). This implies that *S*.Ent induces distinct pathways of cell death via Casp1/11/GSDMD and Casp8. Casp3 cleaves several proteins that mediate apoptosis, including the plasma membrane channel Pannexin-1 (Panx1)^72^. Cleavage and activation of Panx1 leads to formation of a plasma membrane channel that fluxes small molecules such as ATP and promotes early plasma membrane permeability during apoptosis^73,74^. Panx1 is also implicated in macrophage lysis in response to apoptotic stimuli^69,75,76^. Notably, *S*.Ent induced Casp8-dependent Panx1 cleavage in both WT and *Casp1/11*^−/−^ macrophages (Fig. 3d, Extended Data Fig. 2c). Given that loss of Casp8 alone does not affect cytotoxicity (Fig. 2), we reasoned that disruption of blockade of Panx1 alone would be unlikely to robustly impact cytotoxicity in response to *Salmonella* infection. Consistently, WT BMDMs treated with the Panx1 inhibitors probenecid or spironolactone^77,78^, or BMDMs lacking Panx1 had largely WT levels of LDH release (Figs. 3e,f). Notably, treatment of *Casp1/11^−/−^* or *Gsdmd^−/−^* BMDMs with either probenecid or spironolactone, or combined deletion of both GSDMD and Panx1 significantly reduced death relative to loss of GSDMD or Casp1/11 alone (Figs. 3e,f). As we could not simultaneously delete both *Panx1* and *Casp1/11* due to their proximate location on chromosome 9, these data together demonstrate that Panx1 contributes to Casp8-dependent, Casp1/11- and GSDMD-independent cell death in response to *S*.Ent. Consistently, dual loss of Casp8 and GSDMD or Panx1 and GSDMD abrogated release of IL-1β (Extended Data Fig. 2d). Moreover, while loss of GSDMD alone had no effect on *S*.Ent bacterial burdens *in vivo*, and *Panx1*^−/−^ mice actually had slightly, but significantly, reduced levels of bacteria in their tissues, combined loss of both GSDMD and Panx1 resulted in significantly elevated systemic bacterial burdens (Fig. 3g), indicating that pyroptosis and apoptosis function together to promote anti-*Salmonella* innate defense. Altogether, these data demonstrate that *Salmonella enterica* induces late Casp1/11-independent cell death via Casp8- and Panx1-driven apoptosis that mediates anti-*Salmonella* innate defense.

**Fig. 3.**
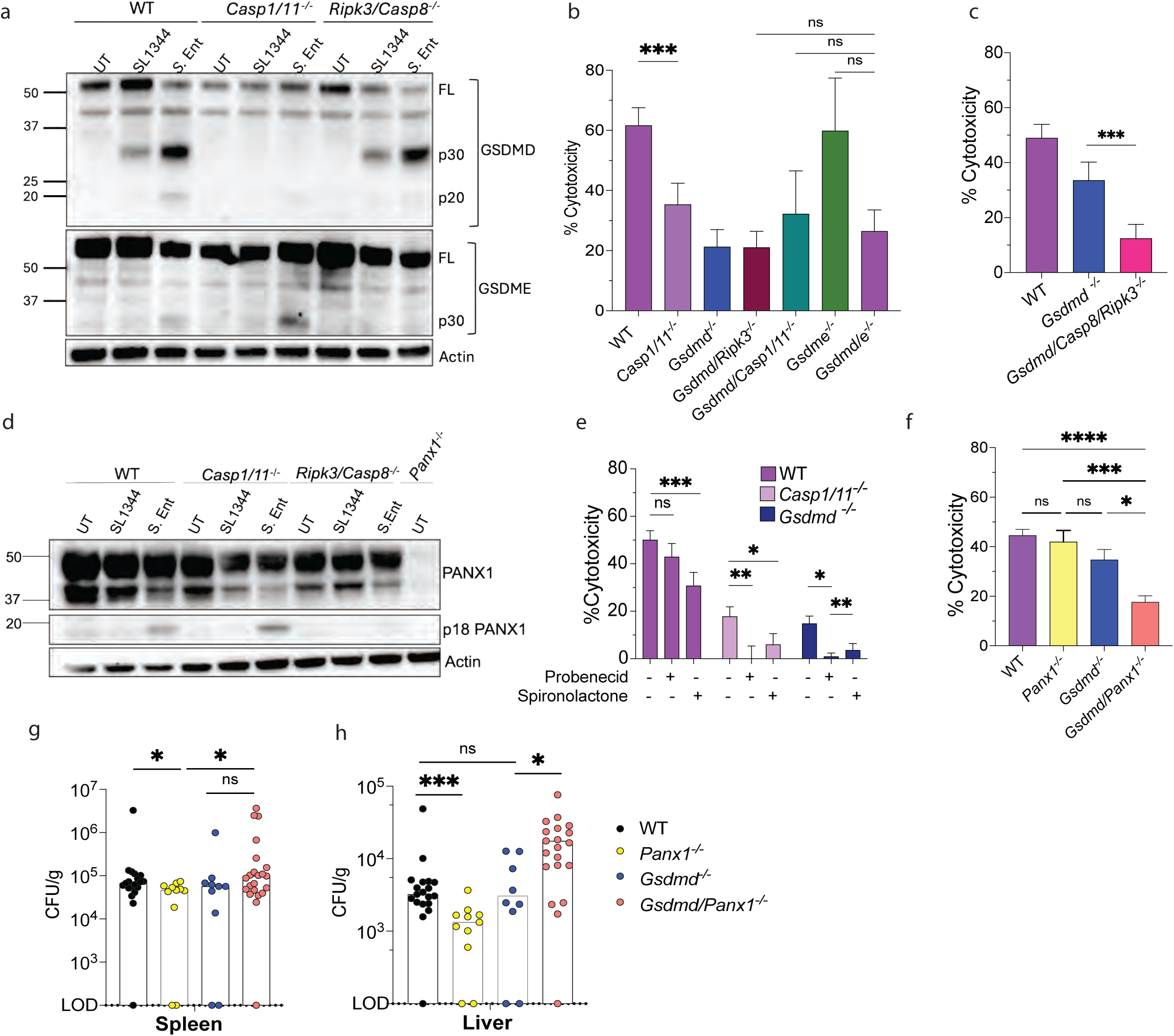
Pannexin-1 promotes Casp8-dependent cell death in response to *Salmonella* infection. **a,** GSDMD and GSDME immunoblots of combined whole cell lysates and supernatants harvested from infected BMDMs 18 hours post-infection with indicated bacterial strains. β-actin as loading control. Data representative of 5 independently performed experiments**. b, c,** Cytotoxicity determined by LDH release in BMDMs infected with *S.*Ent (MOI100) for 18 hours. **d,** Pannexin-1 immunoblot of combined whole cell BMDM lysates and supernatants. **e-f,** Cytotoxicity was determined by LDH after infection with *S.*Ent (MOI100) for 18 hours. **e,** WT, *Casp1/11^−/−^,* and *Gsdmd^−/−^* BMDMs indicated genotypes were treated with either Spironolactone or Probenecid as described in materials and methods. **f,** WT, *Panx1^−/−^*, *Gsdmd^−/−^,* and *Gsdmd/Panx1^−/−^* BMDMs. All bars represent mean ± SEM of at least 3 independent experiments performed in triplicate. \**P*<0.05 \*\**P*<0.01 \*\*\**P*<0.001 \*\*\*\**P<*0.0001 by paired t-test. **g,h,** *S.*Ent bacterial burdens (CFU g^−1^ tissue) in the spleen, liver, and mesenteric lymph-nodes at 15 days post-infection. Mann– Whitney *U* test (two-tailed) for bacterial burdens. \**P* < 0.05, \*\*\**P* < 0.001; NS, not significant.

### Caspase-8-dependent cell death is linked to intracellular *Salmonella* replication

SL1344 is a mouse-virulent derivative of a clinical *Salmonella* Typhimurium isolate, ST4/74, isolated from an infected calf^47,48^, that harbors a mutation in HisG, the first enzyme of the histidine biosynthetic pathway. SL1344 is therefore defective in intracellular replication due to its inability to synthesize histidine^49,79^. While SL1344 and its derivatives have been extensively studied in their interactions with the innate immune system, how histidine auxotrophy and the inability of SL1344 to replicate in primary macrophages affects innate responses to *Salmonella* infection has never been directly investigated. Intriguingly, either exogenous histidine, which restores intracellular replication to SL1344^49,79^, or infection by a prototrophic SL1344 revertant, SL1344^HisG+^, induced increased cell death in WT BMDMs, that was Casp1/11-independent and Casp8-dependent (Fig. 4a, b). Conversely, deletion of *hisG* in *S*.Ent or the histidine prototrophic S.Tm strain, ATCC14028, reduced cell death in WT BMDMs, and abrogated Casp1/11-independent cell death, which was rescued by exogenous histidine (Fig. 4c, Extended Data Fig. 3a). Consistently, in contrast to SL1344, SL1344^HisG+^ induced processing of caspases-8, −3, −7 in both WT and *Casp1/11^−/−^* BMDMs (Fig. 4d). Altogether, these findings demonstrate that Casp1/11-independent, Casp8-dependent cell death is triggered by infection with histidine prototrophic *Salmonella*. As expected, histidine prototrophs, including ATCC14028, as well as histidine auxotrophs supplemented with exogenous histidine, exhibited nearly 10-fold over an 18-hour time-course, suggesting the possibility that Casp8-dependent late apoptosis is triggered by some feature of bacterial replication (Fig. 4e, Extended Data Fig. 3b). Notably, deletion of *hisG* in ATCC14028 abrogated both bacterial replication and Casp1/11-independent cell death, which was restored by histidine supplementation (Extended data Fig. 3a, b). We observed significantly higher numbers of *S.*Ent relative to SL1344 within individual macrophages 12 hours post-infection, corroborating our observations in bulk-replication assays (Fig. 4g, h). We also observed a significant population of highly-infected cells that contained greater than 50 bacteria per cell in *S.*Ent infection that was virtually absent in SL1344-infected cells (Fig. 4i).

**Fig. 4.**
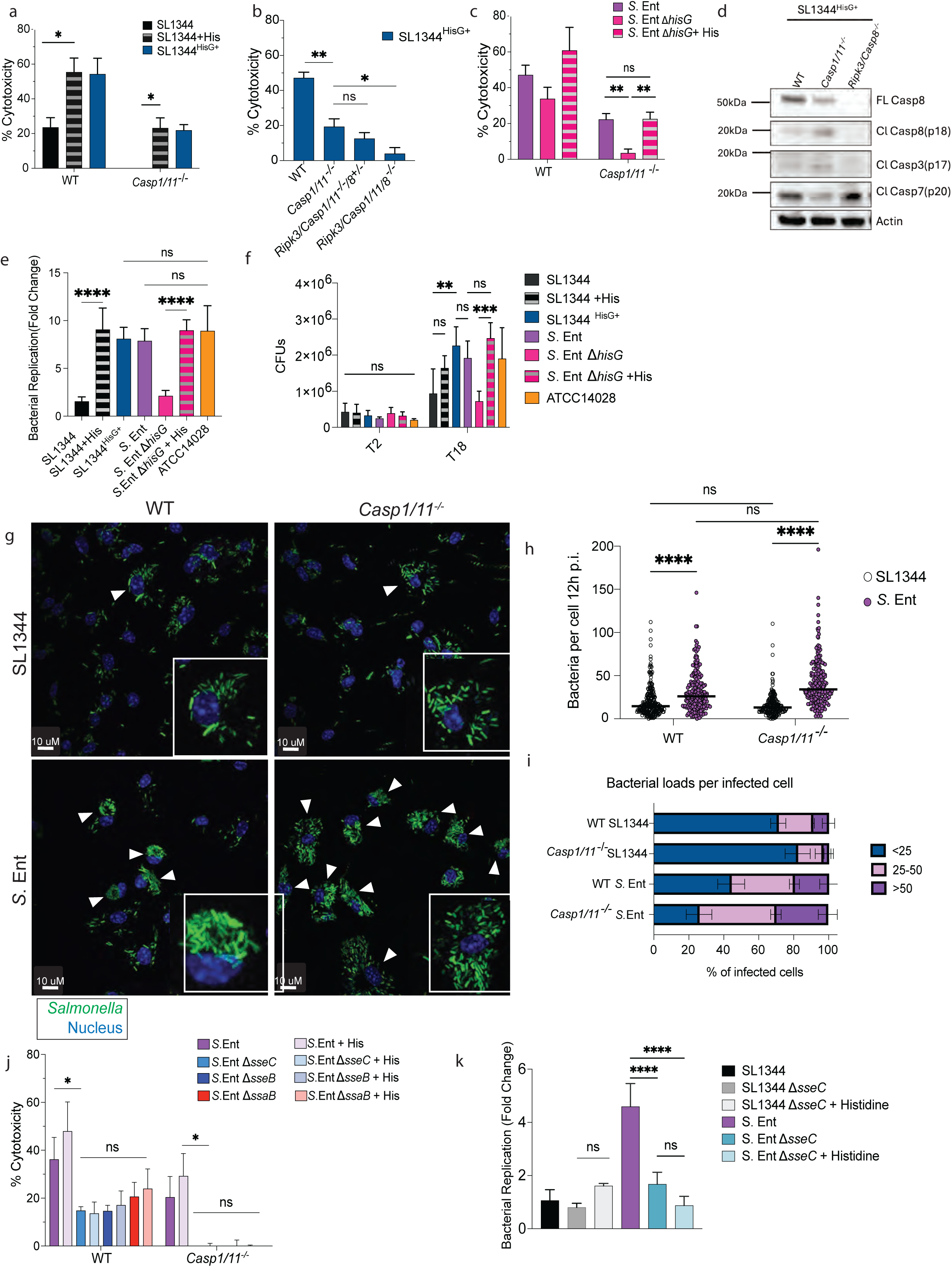
Caspase-8-dependent cell death is linked to intracellular *Salmonella* replication. BMDMs were infected at an MOI of 100 for 18 hours and assayed for cytotoxicity or replication as indicated. **a-c,** Cytotoxicity determined by LDH release in WT and *Casp1/11^−/−^* BMDMs infected with indicated strains. **d,** Combined whole cell lysates and supernatants from indicated BMDM genotypes infected with indicated bacterial strains were analyzed for caspase-8 or −3 cleavage Data representative of 5 experiments. **e,f,** Bacterial replication in *Ripk3/Casp1/11/8^−/−^* BMDMs was determined by CFU present at 18 hours post-infection relative to internalized CFU present at 2 hours post-infection. Mean ± SEM of at least 3 independent triplicate experiments. **g,** WT and *Casp1/11^−/−^* BMDMs were infected for 12 hours with indicated SL1344 and *S.*Ent strains expressing GFP, counterstained with Hoechst dye, and imaged by confocal microscopy. White arrows indicate highly infected (jackpot) cells containing over 50 bacteria per cell. **h,** Quantification of bacteria per infected cell enumerated by manual counting. Data cumulative of 3 independent experiments. **i,** Binning of data in (**h**) classifying low infected cells (n<25), medium infected cells (n=25-30), and the jackpot cells (n>50). **j,** Cytotoxicity determined by LDH release in WT and *Casp1/11^−/−^* BMDMs infected with SPI-2 mutant *S.*Ent. **k,** Bacterial replication in *Ripk3/Casp1/11/8^−/−^* BMDMs was determined by CFU present at 18 hours post-infection relative to internalized CFU present at 2 hours post-infection. Data represent mean of at least three independent experiments each performed in triplicate. \**P*<0.05 \*\**P*<0.01 \*\*\*\**P*< 0.0001 by One-way ANOVA.

Similar to histidine prototrophy, SPI-2 is critical for intracellular bacterial replication in murine BMDMs ^80,81^. Notably, deletion in *S*.Ent of the SPI-2 genes that encode essential translocon components *sseC* and *sseB*, or the SPI-2 effector *ssaB*, which is essential for translocation of other SPI2 effectors^3,80,82^, reduced cytotoxicity in WT BMDMs, and abrogated both Casp1/11- independent cell death and bacterial replication (Fig. 4j, k). The replication defect of SPI-2 mutants could not be restored by exogenous histidine, indicating that metabolic prototrophy and SPI-2 make independent contributions to intracellular replication (Fig. 4j, k). Collectively these data imply that intracellular replication mediates Casp1/11-independent, Casp8-dependent late macrophage cell death.

### SPI-2 needle and rod proteins trigger Casp8-mediated apoptosis in the absence of Casp1/11

How replicating intracellular *Salmonella* trigger Casp8-dependent apoptosis is not clear. Casp8 can be engaged downstream of NLRC4/NAIP-dependent sensing of the SPI-1 T3SS in the absence of Casp1, but deletion of SPI-1 genes did not impact Casp8-dependent cell death, indicating that this pathway is not detecting potentially elevated levels of SPI-1 activity during bacterial replication (Extended data Fig. 4a). SPI-2 activity promotes intracellular replication, making it challenging to distinguish between elevated bacterial burden and levels of SPI-2 T3SS activity. However the transcriptional repressor YdgT limits expression of bacterial horizontally acquired genes^83,84^, and deletion of YdgT enhances SPI-2 expression and secretion without affecting bacterial burdens^83^. Intriguingly, SL1344 Δ*ydgT* triggered elevated cell death in WT BMDMs and triggered Casp8 dependent, Casp1/11-independent cell death without increasing intracellular bacterial burdens (Fig. 5a, b). Moreover, *ydgT* mutation in *S.*Ent also triggered elevated Casp1/11- independent cell death without affecting bacterial replication (Extended Data Fig 4a, b). YdgT controls expression of many SPI-2-related genes, including structural T3SS components^83^. While SPI-2 T3SS structural components evade detection by the NAIP/NLRC4 inflammasome, macrophages primed with TLR ligands have elevated responsiveness to immuno-evasive NAIP ligands, including the SPI-2 inner rod (SsaG) and needle (SsaI) subunits^85,86^. Intriguingly, ectopic delivery of SsaG or SsaI using a heterologous *Yersinia* T3SS by fusing them to the YopE N-terminal secretion signal^85^ (Extended Fig. 4d) triggered robust Casp1/11-independent cell death in primed BMDMs, in contrast to the YopE secretion signal alone, and was accompanied by Casp8 activation and processing of GSDMD to its p20 fragment (Fig. 5c, d). This processing did not occur in WT cells, indicating that in the absence of Casp1/11, Casp8 is activated in response to intracellular delivery of SPI-2 structural components. Importantly, Pam priming also induced Casp1/11- independent macrophage cell death in SL1344 infected macrophages, and this death was abrogated in *Ripk3/Casp1/11/8^−/−^* BMDMs, indicating that priming-induced cell death was mediated by Casp8 (Fig. 5e).

**Fig. 5.**
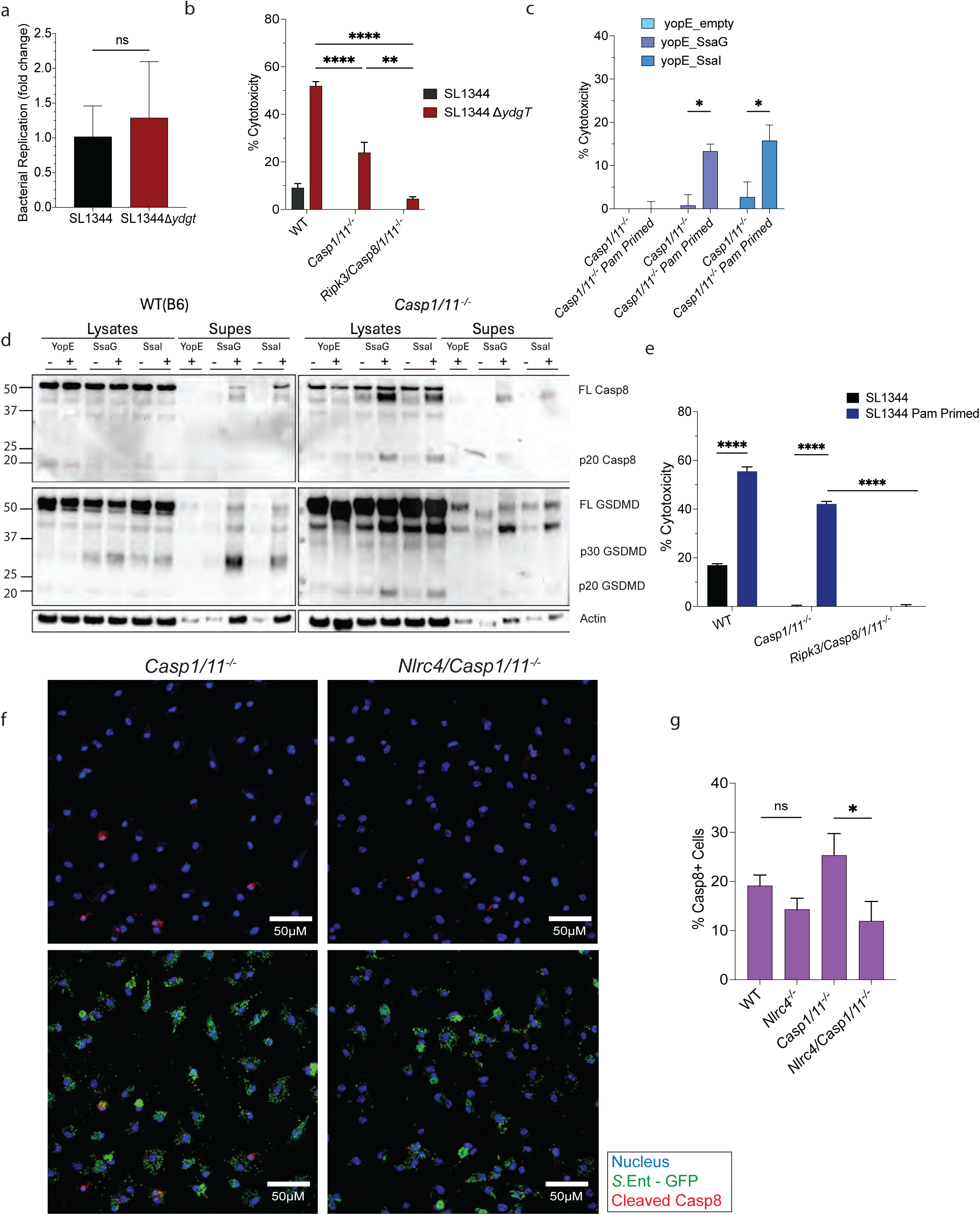
SPI-2 needle and rod proteins trigger Casp8-mediated apoptosis in the absence of Casp1/11. **a,** SL1344 and SL1344Δ*ydgt* replication in *Ripk3/Casp1/11/8^−/−^* BMDMs determined by CFU present at 18 hours post-infection relative to internalized CFU present at 2 hours post-infection. **b,** Cytotoxicity determined by LDH release in WT, *Casp1/11^−/^, Ripk3/Casp8/1/11^−/−^* BMDMs infected with indicated strains BMDMs at MOI of 100 for 18 hours. **c,d,** *Yersinia pseudotuberculosis(Yp)* ΔyopJ expressing either YopE^1–100^ or YopE^1–100^ fused to *S.* Tm SsaG or SsaI used to infect unprimed or Pam3CSK primed WT or *Casp1/11^−/−^* BMDMs at MOI of 20. **c,** Cytotoxicity determined by LDH release 24 h post-infection infected with *Yp* SPI-2 secretion strains for 4 h at an MOI of 20. Cytotoxicity determined by LDH release **d,** Immuno-blot analysis in lysates or supernatants of caspase-8 or GSDMD **e,** Cytotoxicity determined by LDH release in unprimed or Pam3CSK primed WT, *Casp1/11^−/−^*, and *Ripk3/Casp8/1/11^−/−^* BMDMs infected with SL1344. Data cumulative of 3 independent experiments. **f,** *Casp1/11^−/−^, Nlrc4/Casp1/11^−/−^* BMDMs infected for 15 hours with *S.*Ent strains expressing GFP, counterstained with Hoechst dye, and imaged by confocal microscopy. **g,** Quantification of bacteria per infected cell enumerated by manual counting. Data cumulative of 3 independent experiments. \**P*<0.05 \*\**P*<0.01\*\*\**P* < 0.001 \*\*\*\**P*< 0.0001 by paired t-tests.

To test whether in the absence of Casp1/11, NLRC4 was promoting Casp8 activation in response to SPI-2 components from replicating *Salmonella*, we assayed active Casp8 in *S*.Ent-infected *Casp1/11^−/−^* and *Nlrc4/Casp1/11^−/−^* BMDMs. Critically, the frequency of active Casp8-containing BMDMs was significantly elevated in *Casp1/11^−/−^* BMDMs, and this increase was abrogated in *Nlrc4/Casp1/11^−/−^* macrophages (Fig. 5f, g), indicating that the elevated *S*.Ent-induced Casp8 activity in Casp1/11-deficienct BMDMs is NLRC4-dependent. Collectively, these data suggest that in the absence of Casp1/11, *Salmonella* replication leads to increased secretion of SPI-2 T3SS components that trigger NLRC4-dependent Casp8 activation. Notably however, *Nlrc4^−/−^* single knockout cells had similar levels of active Casp8 to WT cells, and loss of NLRC4 and Casp1/11 also did not completely cell death or Casp8 activity, suggesting that Casp8 and Casp8-dependent apoptosis are activated independently of NLRC4 when Casp1 is present, via a pathway that is likely functional even in the absence of NLRC4 and Casp1 (Fig. 5f, Extended Data 4e).

### Replicating *Salmonella* promote caspase-8 activation via TLR4 and TNF signaling

NLRC4-independent Casp8 activation could occur through multiple pathways. While TLRs detect individual PAMPs in a dose-responsive manner, TLR signaling is not thought to be directly linked to detection of bacterial replication per se. Indeed, IL-6 secretion was similar in SL1344- and S.Ent-infected cells (Extended Data Fig. 1i). Unexpectedly however, *S*.Ent-infected BMDMs secreted significantly more tumor necrosis factor (TNF) relative to SL1344-infected BMDMs, and this was dependent on TLR4 (Fig. 6a). SL1344 and *S*.Ent could synthesize different LPS or peptidoglycans that might account for this apparent difference. Critically, using isogenic strains of SL1344 and *S*.Ent, we observed that a histidine prototrophic revertant of SL1344 induced significantly *higher* levels of TNF production, while the histidine auxotrophic *S*.Ent mutant induced significantly *lower* levels of TNF than their isogenic counterparts, demonstrating that TNF secretion is linked to intracellular bacterial burdens (Fig. 6b, c). Intriguingly, TLR4 signaling was required for Casp1/11-independent cell death as well as processing of Casp1 and Casp8 indicating that TLR4 signaling promotes Casp1 activation as well as Casp8-dependent, Casp1/11-independent cell death in response to *Salmonella* replication (Fig. 6d, e). Critically, pre-treatment of BMDMs with either LPS or TNF significantly upregulated Casp1/11-independent cytotoxicity in SL1344-infected cells, which was abrogated in the absence of Casp8, indicating that TLR4- and TNFR1-induced signaling are sufficient to induce Casp8-dependent cell death in response to replicating *Salmonella* (Fig. 6f). Casp8 activation occurs downstream of TNF receptor signaling^37,87,88^. Intriguingly, while we observed robust levels of active Casp8 in *S.*Ent or SL1344^HisG^*-*infected BMDMs, unlike in SL1334-infected BMDMs, Casp8 activation was abrogated in *Tnfr1^−/−^* BMDMs, indicating that Casp1/11-indpendent, Casp8-dependent cell death is mediated by TNFR1 signaling (Fig. 6g, h). Interestingly, while TNFR1 signaling was required for Casp8 activation, inhibition was not enough to abrogate cell death in macrophages unless *Nlrc4* and *Casp1/11* were also absent (Fig. 6i). Altogether, these data reveal that in addition to the well-appreciated pathways of Casp1 and Casp11-mediated pyroptosis triggered by intracellular *S*.Tm, *Salmonella enterica* activates Casp8-dependent apoptosis via engaging TLR4 and TNF signaling downstream of *Salmonella* replication.

**Fig. 6.**
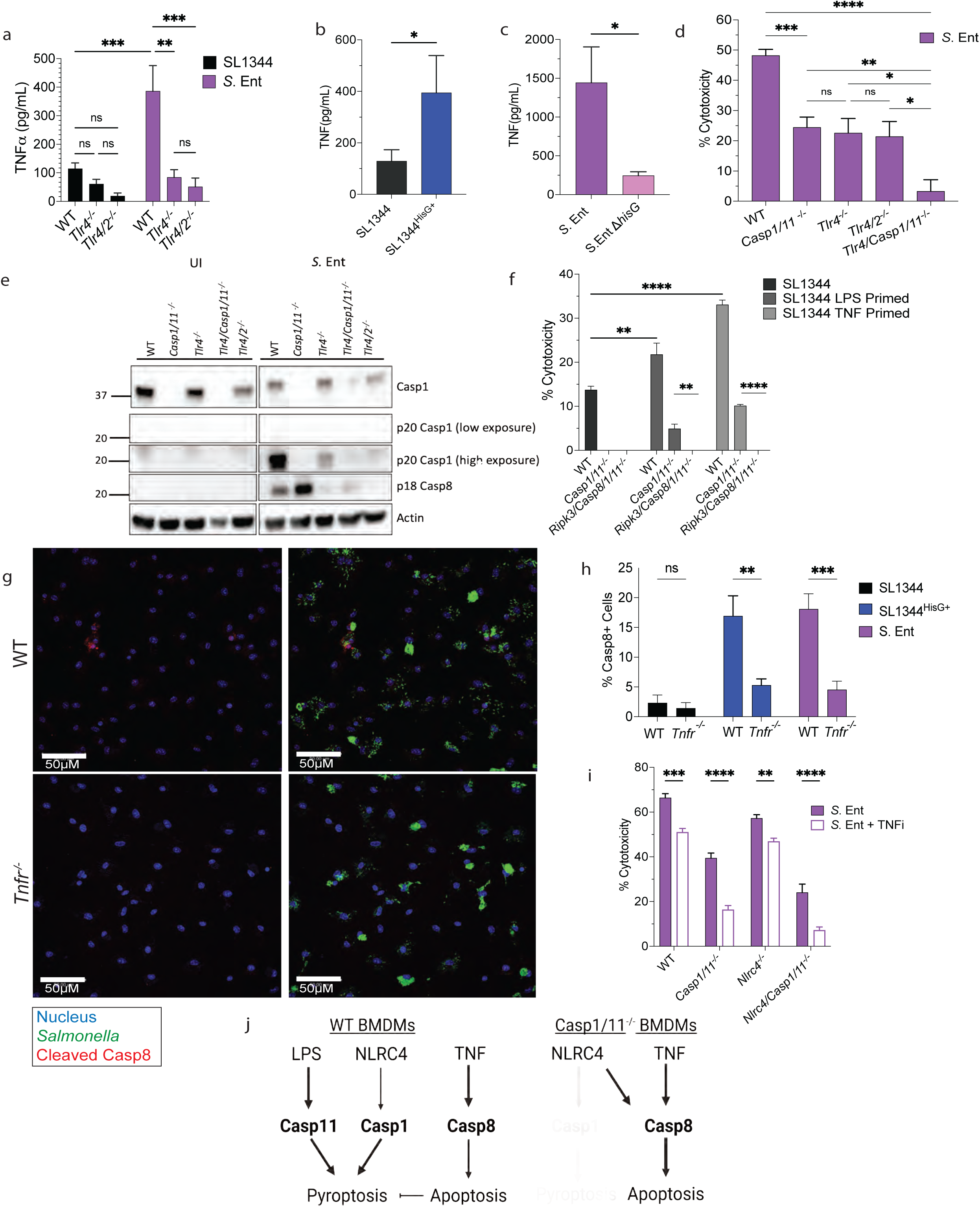
Replicating *Salmonella* promote caspase-8 activation via TLR4 and TNF signaling. **a**,**b,c,** TNFα secretion from BMDMs infected with indicated strains. **d,** Cytotoxicity determined by LDH release in WT, *Tlr4^−/−^, Tlr4/2^−/−^, Tlr4/Casp1/11^−/−^* BMDMs infected with *S.*Ent at MOI of 100 for 18 hours. **e,** Immuno-blot analysis of Casp8 and Casp1 in combined lysates and supernatants of uninfected or *S.*Ent infected BMDMs. **f,** BMDMs were primed overnight with either LPS or TNFα or left unprimed then infected with SL1344. Cytotoxicity determined by LDH. Data represent mean ± SEM of 4 independent experiments. **g,** WT or *Tnfr^−/−^* BMDMs infected for 15 hours with *S.*Ent strains expressing GFP, stained with antibody for cleaved Casp8, counterstained with Hoechst dye, and imaged by confocal microscopy. **h,** Quantification of bacteria per infected cell enumerated by manual counting of BMDMs infected with SL1344, SL1344^HisG+^, or *S.*Ent. Data cumulative of 3 independent experiments. **i,** WT, *Casp1/11^−/−^, Nlrc4^−/−^, Nlrc4/Casp1/11^−/−^* BMDMs were infected with *S.*Ent and either mock treated or treated with TNF inhibitor cocktail. Data represent mean ± SEM of 3 independent experiments. \**P*<0.05 \*\**P*<0.01\*\*\**P* < 0.001 \*\*\*\**P*< 0.0001 by paired t-test. **j,** Model of Casp8 activation in the presence of Casp1/11 and in the absence.

## Discussion

The innate immune system detects microbial threat levels via a series of checkpoints that distinguish between the presence of soluble microbial components at mucosal surfaces and highly pathogenic organisms that penetrate systemic tissues^89–91^. Intracellular viable bacteria are distinguished from soluble microbial products via labile molecules such as short-lived bacterial RNAs (termed Vita-PAMPs), while virulent pathogens are sensed via pathogen-specific activities that disrupt host signaling pathways or deliver microbial products into the host cell cytosol^92,93^. However, pathogens can adopt both replicating and non-replicating states, and the mechanisms that distinguish between replicating and non-replicating pathogens are not clear. Here, we uncover a pathway of Casp8-dependent apoptosis that responds to replicating intracellular bacteria, and is driven by TLR4-dependent TNF signaling. These findings reveal the presence of host mechanisms that sense the intracellular burden of replicating bacterial pathogens and could potentially play a role in distinct immune responses in the context of switches between latency and replication.

Pyroptosis in response to *Salmonella* infection of macrophages is triggered either by rapid NAIP/NLRC4 inflammasome activation of Casp1 in response to cytosolic flagellin^23,25,94^, or delayed non-canonical inflammasome activation triggered by Casp11 detection of cytosolic LPS during the late intracellular stage of infection^26,28^. In the absence of Casp1, cytosolic delivery of bacterial flagellin leads to Casp8 activation by NAIP/NLRC4^33,41,42,44,95^. However, BMDMs are not known to engage Casp8 in response to the late intracellular stage of *Salmonella* infection^18^. Additionally, while Casp8 has been found to be activated in *Salmonella*-infected macrophages in the absence of Casp1/11, in this setting Casp8 plays a back-up role in response to invasive *Salmonella* in in the setting of highly attenuated auxotrophic *Salmonella* strains^33^.

Our findings reveal a previously-unappreciated set of innate responses that detect replicating *Salmonella* and trigger Casp8-dependent and Pannexin-1-mediated apoptosis that promotes host defense. Our data demonstrate that replicating *Salmonella* induce two arms of Casp8 activation that are distinct from previously described responses: one arm acts as a backup pathway to Casp1 and detects SPI-2 T3SS components, while a second arm is triggered by TLR4-dependent TNF signaling and is a non-redundant contributor to innate anti-bacterial responses that promotes macrophage apoptosis at later stages of intracellular infection. SPI-2 T3SS components are thought to evade innate immune detection mediated by the NAIP/NLRC4 inflammasome^25^. However, primed human macrophages respond to the SPI-2 needle protein SsaG via the NAIP-NLRC4 inflammasome^86^, and both SsaG and the SPI-2 rod subunit, SsaI, are detected by primed murine macrophages via TLR-dependent upregulation of NLRC4^85^. Notably, priming with either TLR ligands or TNF significantly increased Casp1/11-independent cytotoxicity in SL1344-infected cells, indicating that TLR and TNF signaling mediate the enhanced Casp8-dependent response to replicating *Salmonella*.

Unlike the findings that pathogen-driven IKK blockade triggers Casp8-mediated activation of GSDMD^63,67^. Casp8-dependent cell death triggered by *Salmonella* replication does not require GSDMD nor GSDME. How Casp8 mediates GSDMD-independent cell death when Casp1 is absent remains unclear, as no specific pore or other lytic mechanism downstream of Casp8 has been defined^33,41,42,44,95^. Notably, the apoptotic channel Pannexin 1 (Panx1), is cleaved downstream of Casp8 activation^74^, and contributes to Casp1/11-independent cell death. Interestingly, while LDH release in response to replicating *Salmonella* was reduced in *Casp1/11^−/−^* cells, we did not observe differences in PI or SYTOX uptake between WT and *Casp1/11^−/−^* cells, indicating that the membrane is permeable to these small dyes during apoptosis. Notably, while Panx1 inhibitors probenecid or spironolactone completely abrogated cell death in either *Gsdmd^−/−^* or *Casp1/11^−/−^* BMDMs, deletion of Panx1 in GSDMD-deficient cells significantly reduced, but did not completely abrogate, cell death relative to *Gsdmd^−/−^* BMDMs, suggesting additional Casp8-dependent pore forming events occur that mediate apoptosis. However, the combined loss of GSDMD and Panx1 led to increased bacterial systemic burdens during acute infection, indicating combined roles for pyroptosis and Casp8-dependent apoptosis in host defense against *Salmonella*.

Collectively, our data unveil a previously undescribed role for Casp8-dependent apoptosis that occurs in response to sensing intracellular *Salmonella enterica* replication, thus revealing a previously underappreciated and generalized feature of innate immune defense against *Salmonella* that may distinguish between latent and replicating bacterial pathogens.

## Acknowledgments

We thank the Brodsky and Shin labs and Dr. Sunny Shin for fruitful discussions. We thank Dr. Denise Monack for sharing of the DT104 strain. We thank the Penn Vet Imaging Core Facility and Gordon Ruthel (RRID: SCR_022764) for use of the Leica Stellaris confocal microscope that was used for microscopy and was funded through NIH grant S10 OD032305-01A1. This work was also supported by NIH Awards R01AI128530 (IEB), R0AI1139102A1 (IEB), R21AI163596A1 (IEB), a BWF Investigator in the Pathogenesis of Infectious Disease Award (IEB), and F31AI161319-3 (BIH).

## Author Contributions

B.I.H, I.E.B. conceived and designed the project. B.I.H., R.G.S, M.Z. performed the experiments. B.I.H. analyzed the data. B.I.H., R.G.S, M.Z. contributed to generation of knock-out strains and genetic manipulation. L.W.P., J.L.R., K.R. contributed mice and discussions. S.C.R, D.M.S. provided valuable strains for the study. B.I.H, I.E.B. wrote the manuscript. B.I.H., D.M.S., L.W.P, S.C.R., I.E.B, helped edit the manuscript. All authors read and approved the manuscript.

## Material and Methods

### Mouse strains

B6, *Nlrc4^−/−^* ^11^*, Casp1*/*Casp11*^−/−^ mice obtained from Jackson Laboratories and subsequently maintained as a breeding line in-house. *Gsdmd*^−/−^ BMDMs were previously described^46^, provided by Russell Vance, and maintained as a breeding line in-house. *Ripk3^−/−^* mice^96^ were a gift of Kim Newton and Vishva Dixit (Genentech) and *Ripk3/Casp8^−/−^* mice^97^ were a gift of Doug Green (St. Jude Children’s Research Hospital). *Ripk3/Casp1/11/8^−/−^* and *Ripk3/Casp1/11^−/−^/Casp8^+/−^* mice were provided by Russell Vance^98^. *Casp11^−/−^* mice were originally generated by Junying Yuan^16^ (Harvard University) and kindly provided by Tiffany Horng (Harvard University)*. Panx1^−/−^* mice were previously described^40^ and provided by Lance Peterson and Kodi Ravachandran (Washington University in St. Louis). *Gsdmd/Casp1/11*^−/−^, *Gsdmd/Casp8/Ripk3*^−/−^, *Gsdmd/Panx1*^−/−^, *Nlrc4/Casp1/11^−/−^* were crossed in-housed and routinely genotyped to maintain stable mice lines. Mice were maintained in a specific pathogen-free facility by University Laboratory Animal Resources (ULAR) staff in compliance with IACUC approved protocols.

### Animal infections

C57BL/6 wild-type, *Ripk3/Casp8^−/−^, Ripk3^−/−^, Casp1/11^−/−^, and RIPK1^KD^* were acquired from the Jackson Laboratory and bred at the University of Pennsylvania. All animals were bred by homozygous mating and housed separately by genotype. Mice of either sex between 8-12 weeks of age were used for all experiments. All animal studies were performed in accordance with University of Pennsylvania IACUC-approved protocols (protocol #804523).

For intraperitoneal infections, mice were infected with 0.5-2 × 10^3^ CFUs of overnight cultures of *Salmonella* diluted in phosphate buffered saline. Mice were fasted overnight and infected with 1 × 10^7^ CFUs of *Salmonella* for oral infection. At indicated times after infection, mice were euthanized, and tissues were collected and weighed in 1 ml sterile PBS, bead homogenized (MP Biomedical), and plated at 10-fold dilutions on LB plates using asceptic technique to determine bacterial burdens (CFU/g tissue). For *S.*Ent infection of animals treated with RIPK1 kinase inhibitor, C57BL6/J mice were placed on a diet of irradiated Purina 5001 control chow or chow containing RIPK1 inhibitor GSK3540547A (GSKʹ547 chow, provided by GSK) 3 days before infection and allowed to eat ad libitum for the duration of the experiment.

### Bacterial strains and growth conditions

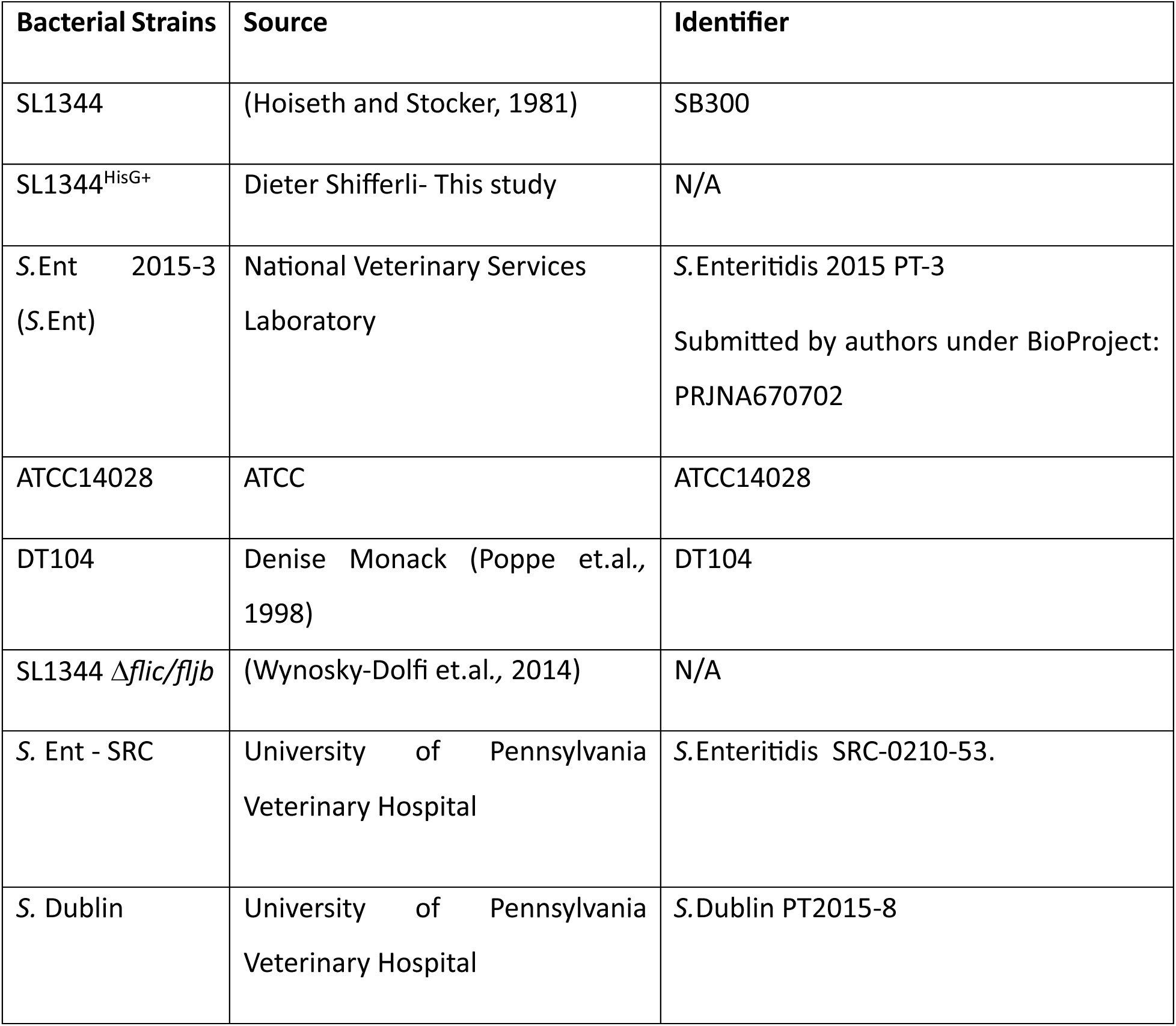

*Salmonella enterica* were routinely grown shaking overnight at 37°C in Luria-Bertani (LB) broth. Targeted deletion strains made in this study were generated in the LT2 strain background of *S. enterica* Typhimurium using standard methods^99^. P22 transduction was used to introduce mutations and antibiotic cassettes into target strains^100^. When necessary, clean deletions were generated using the FRT recombinase as previously described^99^. SL1344 and *S*.Ent fluorescent bacteria harbored pDigC plasmid constitutively expressing GFP^101^.

For SPI-1 induction prior to infection, overnight cultures were diluted in LB with appropriate antibiotics containing 300 mM NaCl and grown standing for 3 hours at 37°C^102^. For non-inducing conditions, strains were grown to stationary phase overnight with aeration for 16-20h at 37°C in LB. For mouse infections, bacteria were grown overnight with aeration at 37°C and diluted in PBS.

### Macrophage infections

Bone-marrow cells harvested from the femur and hips of indicated mouse genotypes were grown at 37°C in a humidified incubator in DMEM containing HEPES, 10% FCS, and 30% L929 supernatant for 7–8 days for differentiation to macrophages. Differentiated bone-marrow derived macrophages (BMDMs) were replated into −12 or 96-well dishes 16–20 h before infection.

For infections examining responses to intracellular *Salmonella,* bacteria were grown to stationary phase overnight for 18-20h. Bacteria were pelleted at 14000 x rpm for 1 minute, washed 2x in DMEM, and resuspended in complete DMEM (cDMEM). BMDMs were infected with *Salmonella* at a multiplicity (MOI) of 100. Infected cells were centrifuged at 1000 × rpm for 5 minutes and incubated at 37°C. 60 minutes post-infection, the cells were treated with 100 *μ*g/mL of gentamicin to kill extracellular *Salmonella*. 60 minutes later, supernatant was removed and replaced with cDMEM containing 10*μ*g/mL of gentamicin. Plates were returned to 37°C for 60 minutes. Media was then replaced with cDMEM containing 10*μ*g/mL of gentamicin for the remainder of the infection. These infections proceeded for 16-19 hours at 37°C as indicated.

For infections examining responses to SPI-1 inducing *Salmonella*, overnight cultures were diluted into LB broth with streptomycin (100μg/mL) containing 300 mM NaCl and grown for 3 hours standing at 37°C to induce SPI-1 gene expression^102^. BMDMs were infected with *Salmonella* at a MOI of 20. Bacterial cultures were then washed in DMEM, pelleted at 14000 × rpm for 1 minute, and resuspended in cDMEM. Infected cells were centrifuged at 1000 × rpm for 5 minutes and incubated at 37°C. 60 minutes post-infection, the cells were treated with 100*μ*g/mL of gentamicin to kill any extracellular *Salmonella*. Where indicated, histidine was supplemented at a final concentration of 250μg/mL. Infections proceeded were allowed to proceed for 2-4 hours at 37°C as indicated. For all experiments, control cells were mock-infected with cDMEM. For priming experiments, BMDMs were primed 3 hours after plating cells and 18 hours prior to infection at the indicated concentrations: 0.5 μg/mL Pam3CSK4 (Invivogen), 100ng/mL LPS(Sigma), 50ng/mL TNFα(Biolegend). For inhibitor experiments, BMDMs were treated either 3 or 6 hours after infection at the indicated concentrations with the following inhibitors: 20μM of the pan-caspase inhibitor Z-VAD(OMe)-FMK (SM Biochemicals; SMFMK001), 30μM GSK‘872 (R&D Systems), 2mM Probenecid (Sigma-Aldrich), 80μM Spironolactone (Sigma-Aldrich), 20μM Nec-1s (Calbiochem), 10μM Necrosulfonamide (Selleck Chemicals), 150μM TAPI-2 (Sigma-Aldrich), and 2mg/ml Ultra-LEAF Purified anti-mouse TNFα antibody (Biolegend). Corresponding solvent for each inhibitor treatment were used as vehicle controls.

### Cell death assays

BMDMs were seeded into 96-well plates at a density of 7 × 10^4^ cells per well. Cells were infected as described above and supernatants harvested at between 2-18 hours after infection as indicated. Lactate dehydrogenase release was quantified using the Cytotox96 Assay kit (Takara Bio, Inc.) according to manufacturer’s instructions and normalized to mock-infected cells. To measure cytotoxicity by uptake of propidium iodide (PI), BMDMs were infected in 96-well black-walled tissue culture plates. At the time when media was replaced with lower gentamicin concentration, 5μM PI was added to plate reader media (20 mM HEPES buffer and 10% FBS in Hank’s Balanced Salt Solution-HBSS). Cells were then allowed to equilibrate to 37°C for 10 minutes before being placed into an IncuCyte live-image analysis system. PI uptake into cells was then measured at an excitation wavelength of 530 nm and an emission wavelength of 617 nm. PI uptake was normalized to mock-infected cells and 1% Triton-treated cells.

### Live Cell Imaging

BMDMs were infected as detailed above. Two hours post-infection, media was removed and exchanges for HBSS containing 10μg/mL of gentamicin and Casp3/7 green reporter (Satori Systems), or Propidium iodide (Sigma Aldrich). Plates were then placed within an IncuCyte imaging system (Sartorius) within an incubator at 37°C, 5% CO2 incubator the remaining 20 hours. Images were taken of the cells every hour at 40x magnification. 4 images were taken per well and cells were enumerated by eye. Percent positivity calculated as total number of PI or Casp3/7 positive cells over total cells in given field.

### Western blotting and antibodies

BMDMs were seeded into 24-well plates at a density of 3 × 10^5^ cells per well and infected with bacteria as described above. 3 h after infection, cells were lysed in 20 mM Hepes, 150 mM NaCl, 10% glycerol, 1% Triton X-100, and 1 mM EDTA. Lysates were mixed with protein loading buffer, boiled, centrifuged, and 20% of the total cell lysate loaded onto 4–12% NuPAGE gels (Invitrogen). Proteins were transferred to PVDF membrane (Millipore). Secreted proteins were isolated from cell supernatants by centrifugation at 2000 rpm for 10 minutes to remove cellular debris, followed by precipitation using trichloroacetic acid (TCA) overnight. Precipitated protein was pelleted by spinning at 13,000 rpm for 15 minutes at 4°C, then washed with ice-cold acetone, centrifuged at 13,000 rpm again for 10 minutes, before finally being suspended in 1x SDS/PAGE sample buffer. Lysates and supernatants were either combined or kept separate as indicated in the figure legend. Primary antibodies specific for cleaved caspase-8 (Asp387/D5B2)(Cell Signaling: 8592S), cleaved caspase-3 (Asp175) (Cell Signaling: 9661S), caspase-8 (Enzo Life sciences: clone IG12 ALX 804-447-C100)(Cell Signaling: D35G2), caspase-3 (Cell Signaling: 96623), caspase-7 (Cell Signaling: 9491T), GSDMD (EPR19828)(abcam: ab209845), GSDME (EPR19859) (abcam: 215191), Pannexin1 (Cell signaling: 00501), IL-1beta (R&D Systems: AF-401-NA), or mouse anti-mouse β-actin (Sigma). Secondary antibodies used were goat anti–rabbit HRP, goat anti-rat HRP (Jackson ImmunoResearch Laboratories, Inc.), goat anti-mouse HRP, and donkey anti-goat (Thermo Scientific). ECL Western Blotting Substrate (Pierce/Thermo Scientific) was used as the HRP substrate for detection.

### Intracellular replication assay

BMDMs that undergo minimal cell death during the 17–20-hour time course of *Salmonella* infection (*Ripk3/Casp1/11/8*^−/−^ or *Ripk3/Casp1/11*^−/−^*/8*^+/−^) were infected with *Salmonella* strains as described above at an MOI of 100. 1-hour post-infection, cells were treated with 100μg/ml of gentamicin for 60 minutes to kill any extracellular bacteria. Media was then replaced with fresh media containing 1μg/ml of gentamicin with or without 250μg/mL of histidine as indicated. At the indicated time points, the infected cells were lysed with PBS containing 0.1% Triton to collect all intracellular *Salmonella*. Harvested bacteria were serially diluted in PBS and spot-plated on LB agar plates. Plates were incubated overnight at 37°C and CFUs were subsequently counted to determine intracellular growth of bacteria.

### Fluorescence microscopy

BMDMs were plated on glass coverslips in a 24-well plate as described above. Cells were either infected with SL1344 or *S.*Ent 2015-3 harboring plasmid pDiGc constitutively expressing GFP^101^ at an MOI of 100 as described above. pDiGc was a gift from Drs. Sophie Helaine & David Holden (Addgene plasmid # 59322; http://n2t.net/addgene:59322; RRID:Addgene_59322). At the indicated timepoints following infection, cells were incubated with Hoechst containing media for 30 minutes. Cells were washed 2 times with PBS and fixed with 4% paraformaldehyde for 10 minutes. Following fixation, cells were mounted on glass slides with Fluoromount-G mounting medium (SouthernBiotech). For detection of cleaved caspase-8, primary antibody rabbit anti-caspase-8 (Asp387/D5B2)(Cell Signaling: 8592S) and secondary goat-anti rabbit Alexa-Fluor 647 Plus (Thermo Scientific) were used. Coverslips were imaged on an inverted fluorescence microscope (Leica Stellaris 8 Falcon (Grant Opportunity Number PAR-22-079 or Leica SP5/FLIM; Olympus) at a magnification of 60x or 100× as indicated. All images were analyzed using ImageJ (Fiji) software.

### ELISA

Harvested supernatants from infected murine macrophages were assayed for cytokine levels using ELISA kits in accordance with manufacturer’s instructions for murine IL-1α (R&D Systems), IL-6 (R&D Systems), IL-1β (BD Biosciences), and TNF-α (R&D Systems).

### Cytometric Bead Array (CBA)

Harvested supernatants from infected murine macrophages were diluted at various concentrations and assayed for cytokine levels using a cytometric bead array (BD Biosciences) for indicated cytokines (IL-1α and IL-1β).

### *Yersinia*-Mediated Delivery of NAIP Ligands

*Yersinia* strains were obtained in house and generated as previously described^85^. Briefly, the coding region for the terminal 35 amino acids in the D0 domain of respective genes was inserted downstream of the *yopE* promoter sequence and the coding sequence of the first 100 amino acids of *yopE* from *Yp* 32777. Constructs were ligated into expression vector pACYC184 using BamHI and SalI restriction sites. Point mutations of the pACYC184 YopE-*S*. Tm or EPEC FliCD0 constructs were generated with the Q5 site-directed mutagenesis kit (New England Biolabs) following manufacturer recommendations. All constructs were validated by Sanger sequencing.

### Statistical analysis

All cell death data are presented as mean ± S.E.M of indicated number of independent wells in a representative experiment. Mouse data are presented as mean ± S.E.M. of the indicated *n* values. All immunoblots were repeated at least 3 times independently with similar results. Plotting of data and statistical analysis were performed using Prism 10.0 (GraphPad Software) and statistical significance determined using the appropriate test as indicated in each figure legend. One-way analysis of variance (ANOVA) with post hoc Dunnett’s test for comparisons among multiple groups with a single control, or two-way ANOVA with post hoc Tukey test for comparisons among different groups, or Student’s paired*-t-* test for comparison between 2 points with a single group. The significance of in vivo survival data was determined using two-sided log-rank (Mantel–Cox) test. Differences were considered statistically significant if the *P* value less was <0.05.

---

**Extended Data Fig. 1. Multiple *Salmonella* serovars and strains trigger Casp8 dependent cell death independently of flagellin.**

**a,** WT, *Casp1/11^−/−^, Ripk3^−/−^* BMDMs infected with SL1344 or *S.*Ent for 18 hours. **b,** WT or *Casp1/11^−/−^* either mock treated or treated with RIPK3 inhibitor GSK872 and infected with SL1344 or *S.*Ent. **c,** Cytotoxicity in BMDMs infected with non-induced *S*Tm strains ATCC14028 or DT104, *S.*Dublin strain PT2015-8, and *S.*Ent strain SRC (all MOI 100) **d,** Cytotoxicity in BMDMs infected with strains grown under SPI-1 inducing conditions at MOI of 20 **e,** Cytotoxicity 18 hours post-infection with non-induced SL1344, *S.*Ent, and respective flagellin mutants (MOI 100). **f,** Infected cells were stained with Sytox Green, placed in a temperature-controlled plate reader at 37°C, and Sytox uptake measured every 30 minutes. Statistical comparisons between cell death kinetics of SL1344 and *S.*Ent and indicated genotypes. **g,** Immunoblot of full length and processed IL-1β of combined supernatants and whole cell lysates of primary BMDMs infected with SL1344 or *S.*Ent. β-actin as loading control. Data representative of at least 4 experiments. **h,** IL-1α cytokine release from BMDMs determined by Cytometric Bead Array. **i,** IL-6 cytokine release from BMDMs determined by ELISA. Data represent mean ± SEM of at least 3 independent experiments. \**P*<0.05 \*\**P*<0.01 \*\*\**P* < 0.001 \*\*\*\**P* < 0.0001 **a-f,** by One-Way ANOVA **h,i** by Two-Way ANOVA

**Extended Data Fig. 2. Casp8 activation of Pannexin1 permits Casp1-independent IL-1 release.**

**a-c,** Immunoblots of processed **a,** GSDMD **b,** GSDME **c,** Panx1 in combined whole cell lysates and supernatants infected for 18hours. β-actin as loading control. Data representative of at least 5 experiments. **c,** IKK inhibitor and LPS stimulation used as positive control for apoptosis. **d,** IL-1β cytokine release from *S.*Ent infected BMDMs determined by ELISA. Data represent mean ± SEM of at least 3 independent experiments. \**P*<0.05 \*\**P*<0.01 \*\*\**P*<0.001 by One-Way ANOVA

**Extended Data Fig. 3. *Salmonella* strains deficient in histidine metabolism fail to grow within macrophages and fail to trigger Casp8 dependent cell death.**

**a,** WT or *Casp1/11^−/−^* BMDMs infected with ATCC14028 or its isogenic knockout *hisG* strain (MOI100) either mock treated or treated with histidine. Cytotoxicity was determined by LDH release 18 hours post-infection. Data represent mean ± SEM of at least 3 independent experiments**. \****P*<0.05 \*\*\**P*<0.001 by One-Way ANOVA. **b.** Replication of isogenic wild-type ATCC14028 and *ΔhisG* strains determined by CFU present at 18 hours post-infection relative to internatlized CFU at 2 hours post-infection. **c.** Confocal fluorescence microscopy was performed on bone marrow-derived macrophages from indicated genotypes 12 hours post-infection with GFP-expressing SL1344^HisG+^ bacteria.

**Extended Data Fig. 4. SPI-2 needle and rod proteins trigger Casp8-mediated apoptosis in the absence of Casp1/11**

**a,** Cytotoxicity determined by LDH release 18 hours post-infection in WT and *Casp1/11^−/−^* infected with *S.*Ent SPI-1 mutants. Data represent mean ± SEM of at least 3 independent experiments. **b,** Cytotoxicity determined by LDH release 18 hours post-infection in WT and *Casp1/11^−/−^* infected with *S.*Ent or *S.*EntΔ*ydgt* **c,** *S.*Ent and *S.*EntΔ*ydgt* replication in *Ripk3/Casp1/11/8^−/−^* BMDMs determined by CFU present at 18 hours post-infection relative to internalized CFU present at 2 hours post-infection. **d,** Schematic of *Yersinia* constructs harboring SPI-2 SsaG and SsaI. **e,** Cytotoxicity determined by LDH release 18 hours post-infection in WT, *Nlrc4^−/−^, Casp1/11^−/−^,* or *Nlrc4/Casp1/11^−/−^* infected with SL1344, SL1344^HisG+^, or *S.*Ent. Data represent mean ± SEM of at least 3 independent experiments**. \****P*<0.05 \*\**P*<0.01 \*\*\**P*<0.001 by **a,c,e** One-Way ANOVA or **b,** paired t-test.

