## Supplementary figures and images for "Innate detection of *Salmonella* replication triggers caspase-8-dependent apoptosis via TLR-driven TNF signaling and NLRC4-mediated sensing of the SPI-2 Type III secretion system"

### Extended Figures

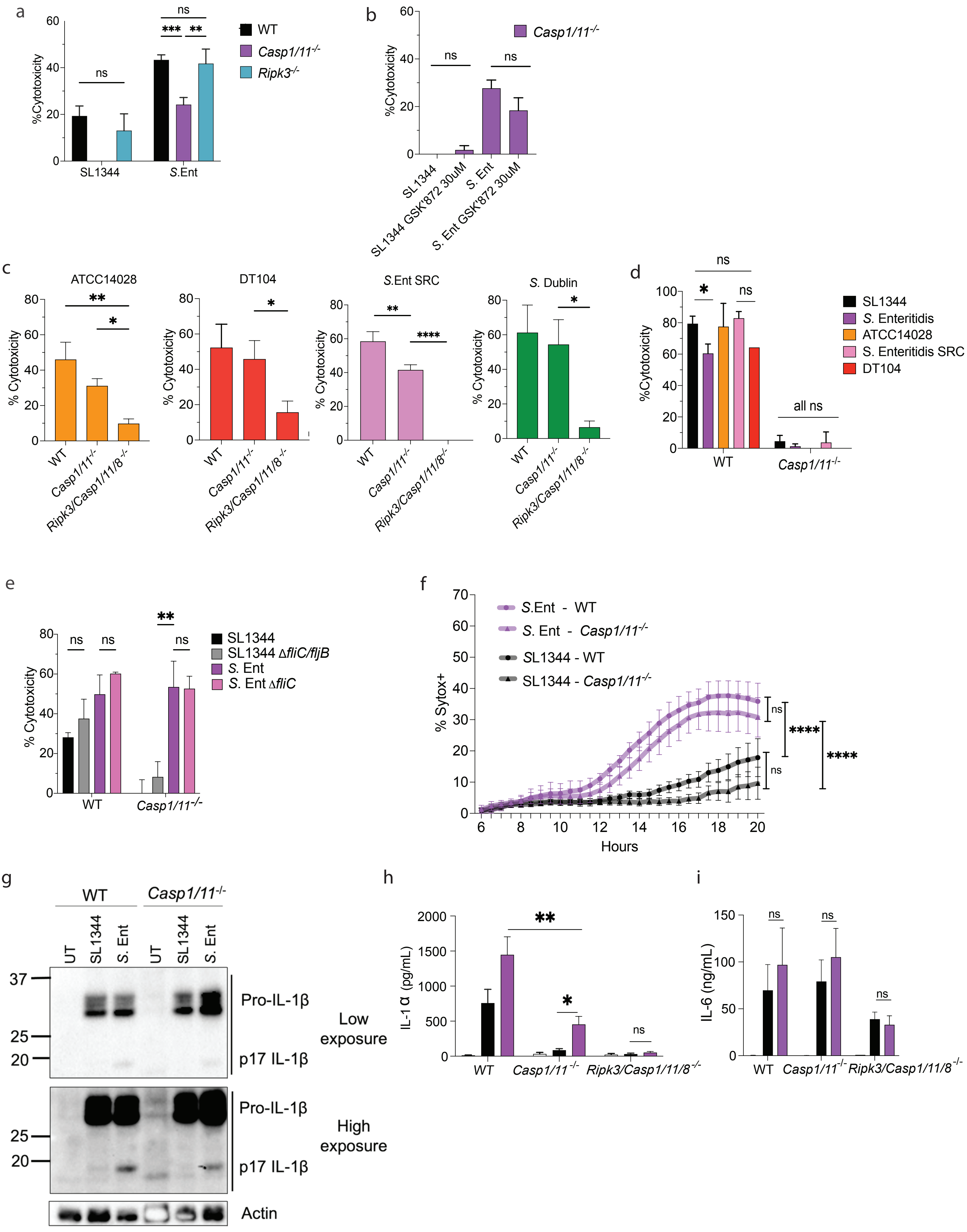


Extended Data Figure 1

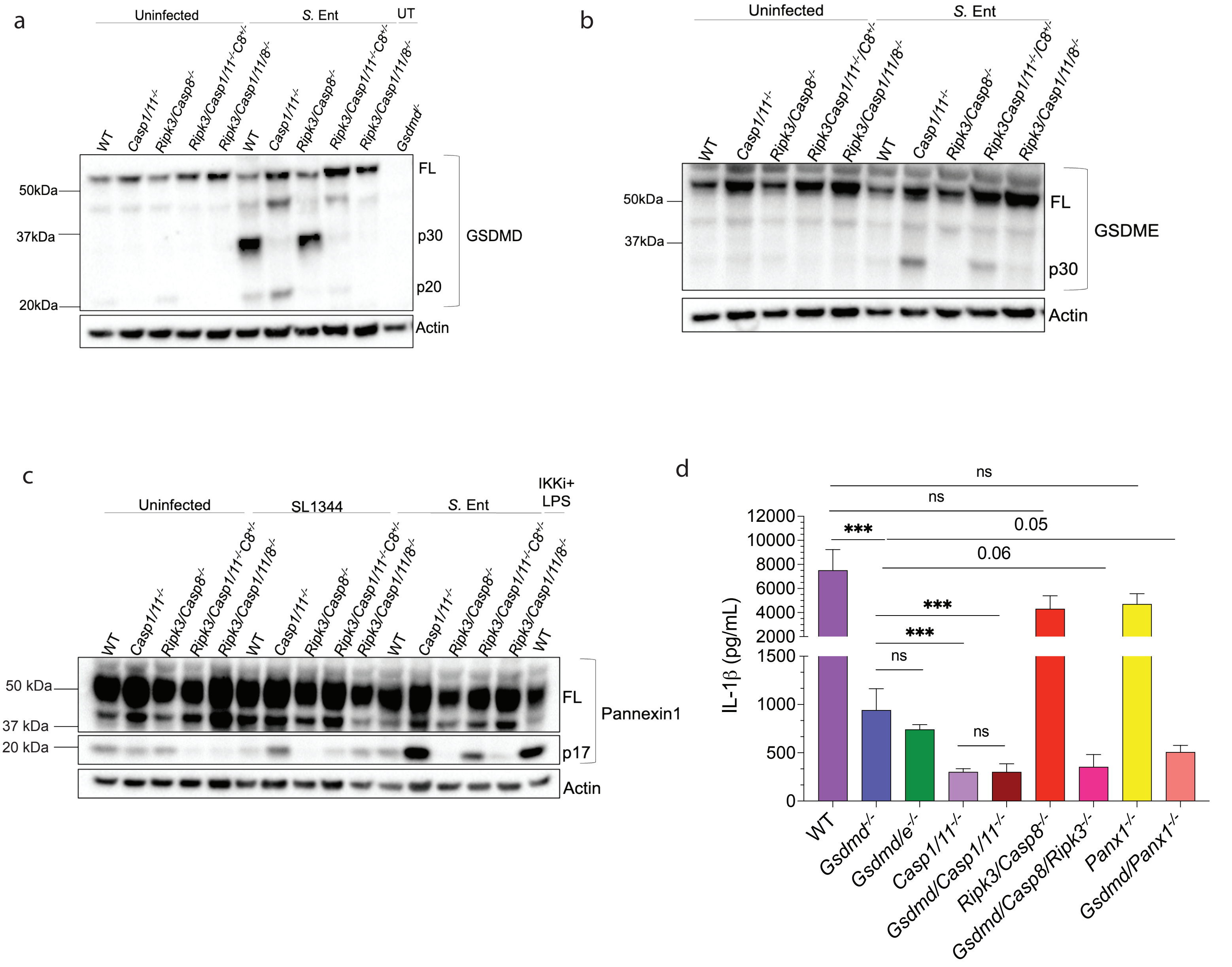


Extended Data Figure 2

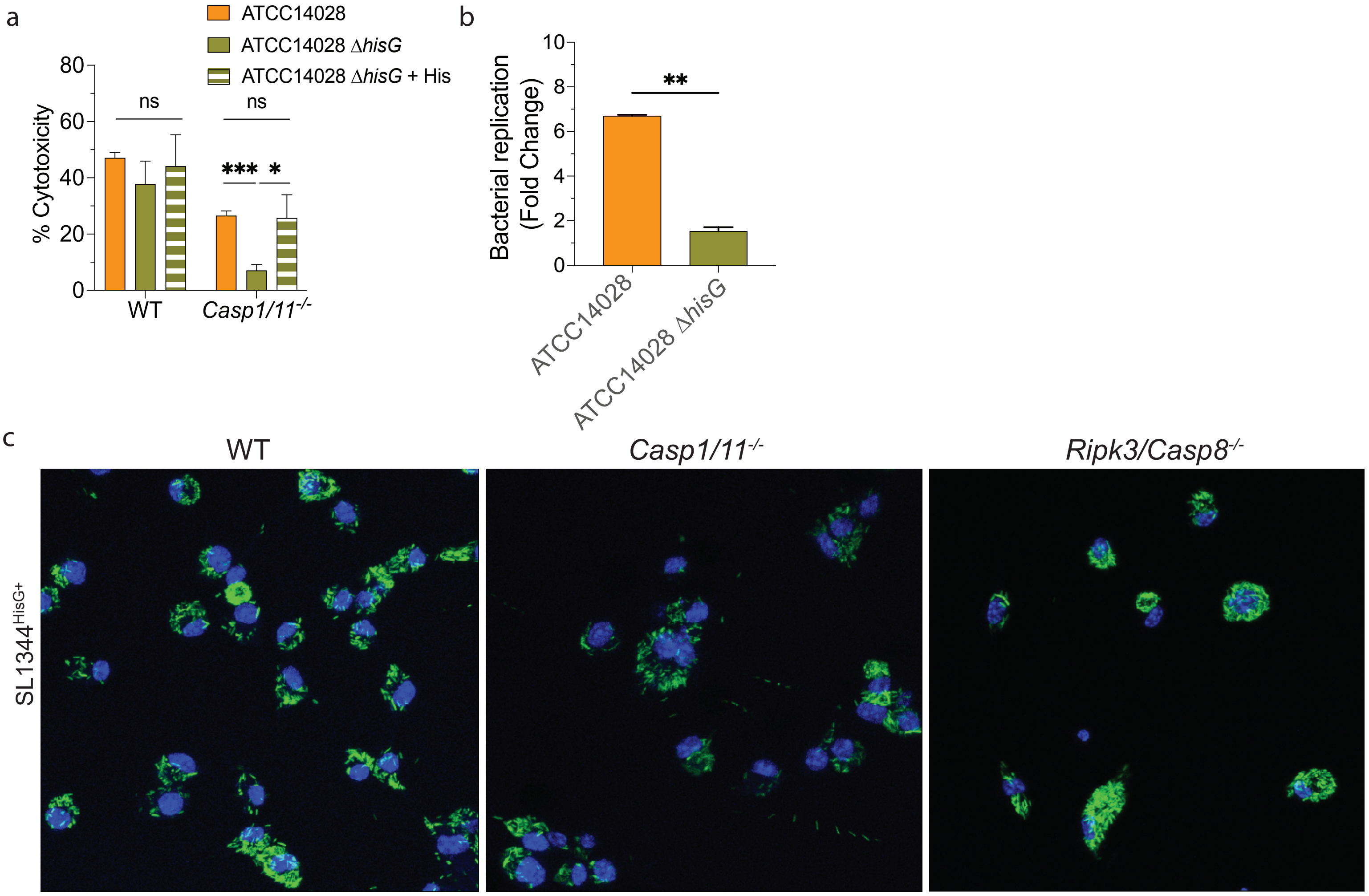


Extended Data Figure 3

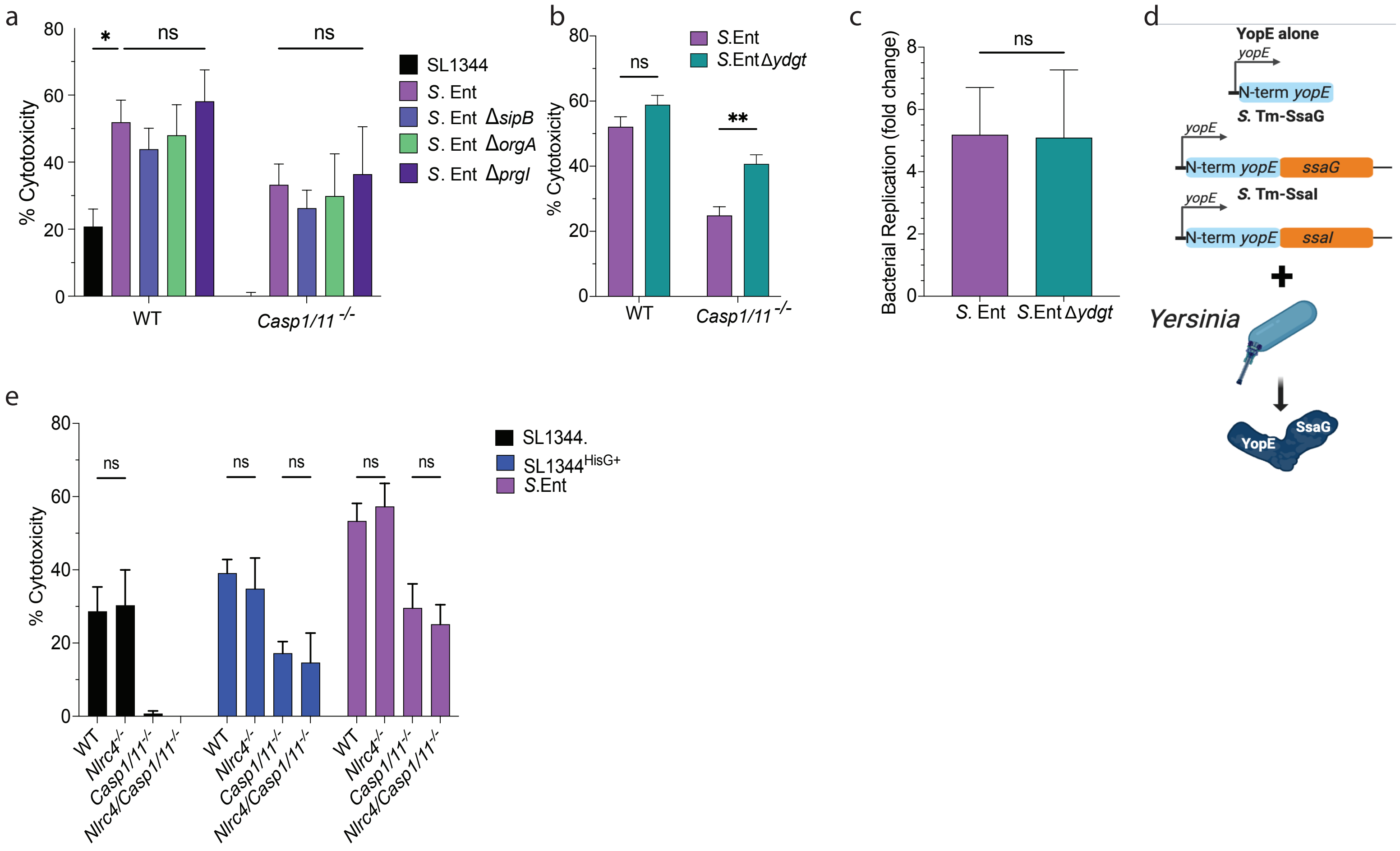


Extended Data Fig 4
